# The *Eruca sativa* genome and transcriptome: A targeted analysis of sulfur metabolism and glucosinolate biosynthesis pre and postharvest

**DOI:** 10.1101/2019.12.23.886937

**Authors:** Luke Bell, Martin Chadwick, Manik Puranik, Richard Tudor, Lisa Methven, Sue Kennedy, Carol Wagstaff

**Affiliations:** School of Agriculture, Policy & Development, PO Box 237, University of Reading, Whiteknights, Reading, Berkshire. RG6 6AR. UK; School of Chemistry Food & Pharmacy, PO Box 226, University of Reading, Whiteknights, Reading, Berkshire. RG6 6AP. UK; Elsoms Seeds Ltd., Pinchbeck Road, Spalding, Lincolnshire. PE11 1QG. UK

## Abstract

Rocket (*Eruca sativa*) is a source of health-related metabolites called glucosinolates (GSLs) and isothiocyanates (ITCs) but little is known of the genetic and transcriptomic mechanisms responsible for regulating pre and postharvest accumulations. We present the first *de novo* reference genome assembly and annotation, with ontogenic and postharvest transcriptome data relating to sulfur assimilation, transport, and utilization. Diverse gene expression patterns related to sulfur metabolism and GSL biosynthesis are present between inbred lines of rocket. A clear pattern of differential expression determines GSL abundance and the formation of hydrolysis products. One breeding line sustained GSL accumulation and hydrolysis product formation throughout storage. Copies of *MYB28*, *SLIM1, SDI1* and *ESM1* orthologs have increased and differential expression postharvest, and are associated with GSLs and hydrolysis product formation. Two glucosinolate transporter gene orthologs (*GTR2*) were found to be associated with increased GSL accumulations.

## Introduction

Sulfur (S) is a critical macronutrient that plants require for growth and development ^1^. Sulfate (SO4^2-^) is utilized as a primary means of synthesizing numerous S-containing metabolites, such as amino acids (cysteine and methionine), glutathione (GSH), and glucosinolates (GSLs) ^2^. GSL compounds are present in species of the order Brassicales, and are abundant in many vegetables and condiments worldwide, such as rapeseed (*Brassica napus*), Chinese cabbage (*Brassica rapa*), cabbage (*Brassica oleracea* var. *capitata*), and broccoli (*B. oleracea* var. *italica*) ^3^. GSLs are also found in the leafy vegetable *Eruca sativa* (“salad” rocket), which has gained significant popularity amongst consumers over the last ten years ^4^. Rocket salad is known for its distinctive flavour, aroma, and pungency, and can be eaten raw without the need for cooking ^5^.

GSLs are synthesised as part of plant defense mechanisms against pests and diseases ^6^, and can also act as important S storage molecules ^1^. Compounds such as glucosativin (4-mercaptobutyl GSL; GSV) and glucorucolamine (4-cystein-S-yl)butyl GSL; GRL) are unique to the genera *Eruca* and *Diplotaxis* (‘wild’ rocket) ^7^. GSV can exist in a dimer form (dimeric 4-mercaptobutyl GSL; DMB), and diglucothiobeinin (4-(β-D-glucopyranosyldisulfanyl)butyl GSL; DGTB) is a unique GSL dimer of these species ^8^.

Aliphatic GSLs are regulated by *MYB28*, *MYB29*, and *MYB76* transcription factors (TFs), and indolic GSLs by *MYB34*, *MYB51*, and *MYB122* ^2^. These MYBs are in turn regulated by basic helix-loop-helix (bHLH) transcription factors such as *MYC2*, which are involved in plant defense response ^9^. Other transcriptional regulators, such as *SLIM1* (*SULFUR LIMITATION 1*) and *SDI1* (*SULFUR DEFICIENCY INDUCED 1*) also interact with MYB transcription factors to regulate the use and efficiency of sulfur within the plant. As GSLs contain significant amounts of sulfur (up to 30% of total plant S-content) the synthesis and catalysis of these compounds is crucial in times of stress (Figure 1) ^10, 11^.

**Fig. 1.**
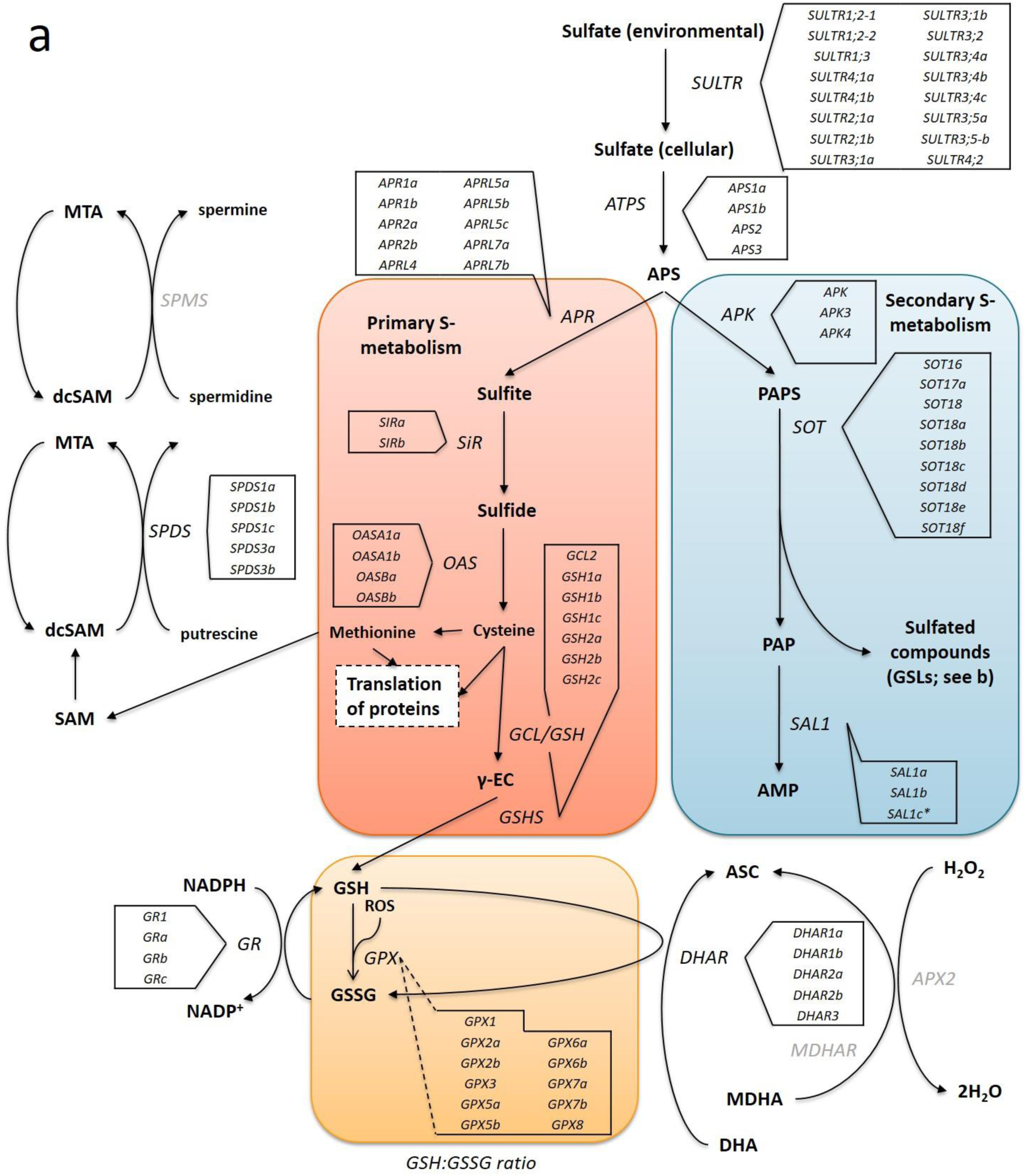
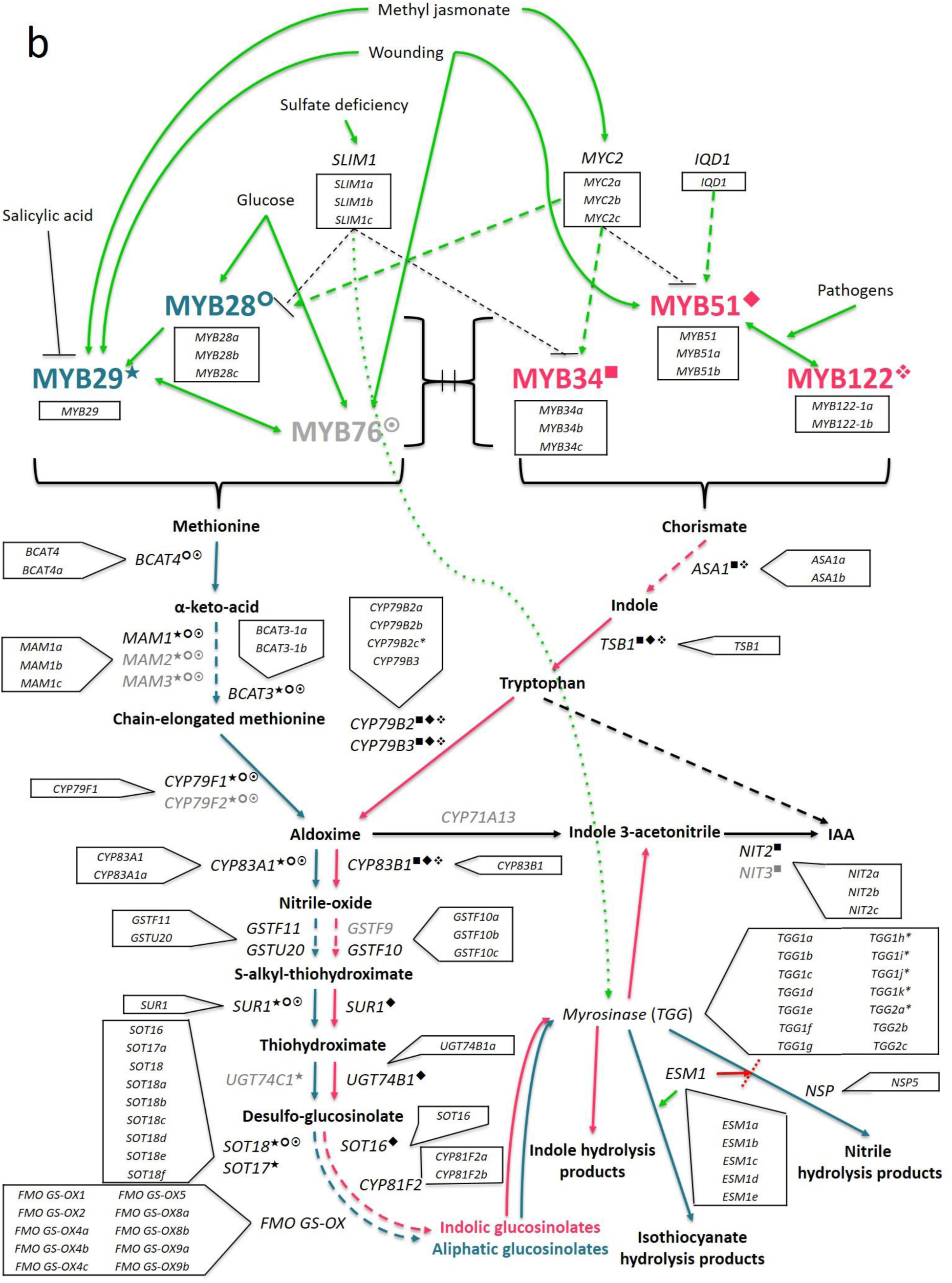
Sulfur metabolism and glucosinolate biosynthesis pathways. The primary (red box) and secondary (blue box) sulfur metabolism pathways (**a**) of *Arabidopsis thaliana* with identified homologous genes within the *Eruca sativa* genome annotation (see boxed insets) adapted from Chan et al. ^11^. Environmental sulfur is assimilated and integrated into key amino acids (cysteine and methionine) and enzymes. Sulfur metabolism is also intrinsically linked with oxidative stress via glutathione synthesis. Under stress conditions 5’-phosphoadenosine-3’-phosphate (PAP), glutathione disulfide (GSSG), and reduced glutathione (GSH) direct sulfate towards GSH production. The GSH:GSSG redox state ratio is also known to influence sulfur assimilation rates (orange box). *SOT* (*sulfotransferase*) genes link secondary sulfur metabolism with the final sulfation step of GSL biosynthesis, and it is thought that *SAL1* plays an important role in regulating the activity of these genes through interaction with PAP. GSL biosynthesis (**b**; adapted from Gigolashvili et al. ^17^) is initiated by a complex and interacting network of abiotic and biotic factors. Aliphatic synthesis pathway shown in teal, and the indolic pathway shown in pink, is regulated by R2R3-MYB transcription factors. Known interactions between MYBs and specific genes within each respective pathway are highlighted as follows: ❍ = MYB28, ★ = MYB29, ⊙ = MYB76, ■ = MYB34, ◆ = MYB51, ❖ = MYB122. Genes with identified orthologs in the *E. sativa* genome annotation are written in black; those with no identified homologous sequence are written in grey. Abbreviations: *SULTR*, *sulfate transporter*; ATP, adenosine triphosphate; *ATPS*, *ATP sulfurylase*; *APR, APS reductase*; *APK, APS kinase*; *SiR, sulfite reductase*; *OASTL, O-acetylserine lyase*; *GCL, glutamate cysteine ligase*; γ-EC, γ-glutamyl-cysteine; *GSHS, GSH synthetase*; *GR, glutathione reductase*; *GPX, glutathione peroxidase*; ASC, ascorbate; DHA, dehydroascorbate; *DHAR, DHA reductase*; MDHA, monodehydroascorbate; *MDHAR, MDHA reductase*; *APX2, ascorbate peroxidase 2*; H2O2, hydrogen peroxide; PAPS, 5’-phosphoadenosine-3’-phosphosulfate; SAM, S-adenosyl methionine; dcSAM, decarboxylated SAM; MTA, methylthioadenosine; *SPDS, spermidine synthase*; *SPMS, spermine synthase*; *SLIM1, sulfur limitation 1*; *IQD1, protein IQ domain 1*; *BCAT, methionine aminotransferase*; *MAM, methylthioalkylmalate synthase*; *CYP, cytochrome P450*; *GST, glutathione-S-transferase*; *SUR1, C-S lyase 1*; *UGT, UDP-glycosyltransferase*; *FMO GS-OX, flavin-containing monooxygenase*; *ASA1, anthranilate synthase alpha subunit 1*; *TSB1, tryptophan synthase beta chain 1*; IAA, indole-3-acetic acid; *NIT, nitrilase*; *ESM1, epithiospecifier modifier protein 1*; *NSP, nitrile specifier protein*.

GSLs themselves are not bioactive, and are hydrolysed by myrosinase enzymes (TGGs) when tissue damage takes place. They form numerous breakdown products including isothiocyanates (ITCs), which are of foremost interest for their anticarcinogenic effects in humans ^12^. Salad rocket produces the ITC sulforaphane (SF; a breakdown product of glucoraphanin; 4-methylsulfinylbutyl GSL, GRA), which has been well documented for its potent anticarcinogenic properties ^13^. SF is abundant in broccoli, however its hydrolysis from GRA is often inhibited or prevented due to high cooking temperatures employed by consumers, which denatures myrosinase at temperatures >65°C ^14^.

A previous study by Bell et al. ^15^ observed that both GSL and ITC concentrations increased significantly in rocket salad post-processing, but that this varied according to cultivar. The authors proposed that in response to the harvesting and washing process, stress responses within leaf tissues were initiated, leading to the increase in synthesis of GSLs and subsequent hydrolysis into ITCs. Sugar content, by comparison, showed little dynamic change and little reduction in the same samples, which could have implications for sensory perceptions and consumer acceptance ^5^.

We present a *de novo E. sativa* reference genome sequence, and report on the specific effects harvest, wash treatment, and postharvest storage have on GSL biosynthesis and sulfur metabolism gene expression through RNA sequencing (RNAseq) in three elite inbred lines. We also present evidence of transcriptomic changes between first and second cuts of rocket plants, and how this in turn leads to elevated concentrations of both GSLs and ITCs. We hypothesised that each rocket line would vary in their ability to retain and synthesize GSLs post washing and during shelf life cold storage, as well as vary in their relative abundances between first and second cuts.

## Results

### *E. sativa* genome assembly and annotation

*De novo* reference genome sequence was produced by interleaving Illumina MiSeq and HiSeq2500 sequence data (Illumina Inc., San Diego, CA, USA). An elite breeding line (designated **C**) was selected for PCR-free paired-end sequencing and long mate-pair end sequencing and assembled into 49,933 contigs (>= 500 bp). The resulting assembly was ∼851 Mb in size (Supplementary Table S1).

Transposable elements (TEs) within the *E. sativa* genome comprise 66.3% of its content. The majority of TEs are long terminal repeat (LTR) retrotransposons (37.3%), with long interspersed nuclear elements (LINEs; 3.3%) and short interspersed nuclear elements (SINEs; 0.3%) having lower relative abundance. 18.2% of all TEs identified were of unknown classification (Supplementary Table S2).

A total of 45,438 protein-coding genes were identified within the assembly, with an average length of 1,889.6 bp, and an average of 4.76 exons per gene. This genome size is smaller than that predicted for radish (*Raphanus sativus*), and larger than *Arabidopsis lyrata* (Supplementary Table S3), and is consistent with what is known of Brassicales phylogeny ^16^. 98.3% of predicted genes were found to have homology with other plant species (Figure 2b, Supplementary Table S4).

**Figure 2.**
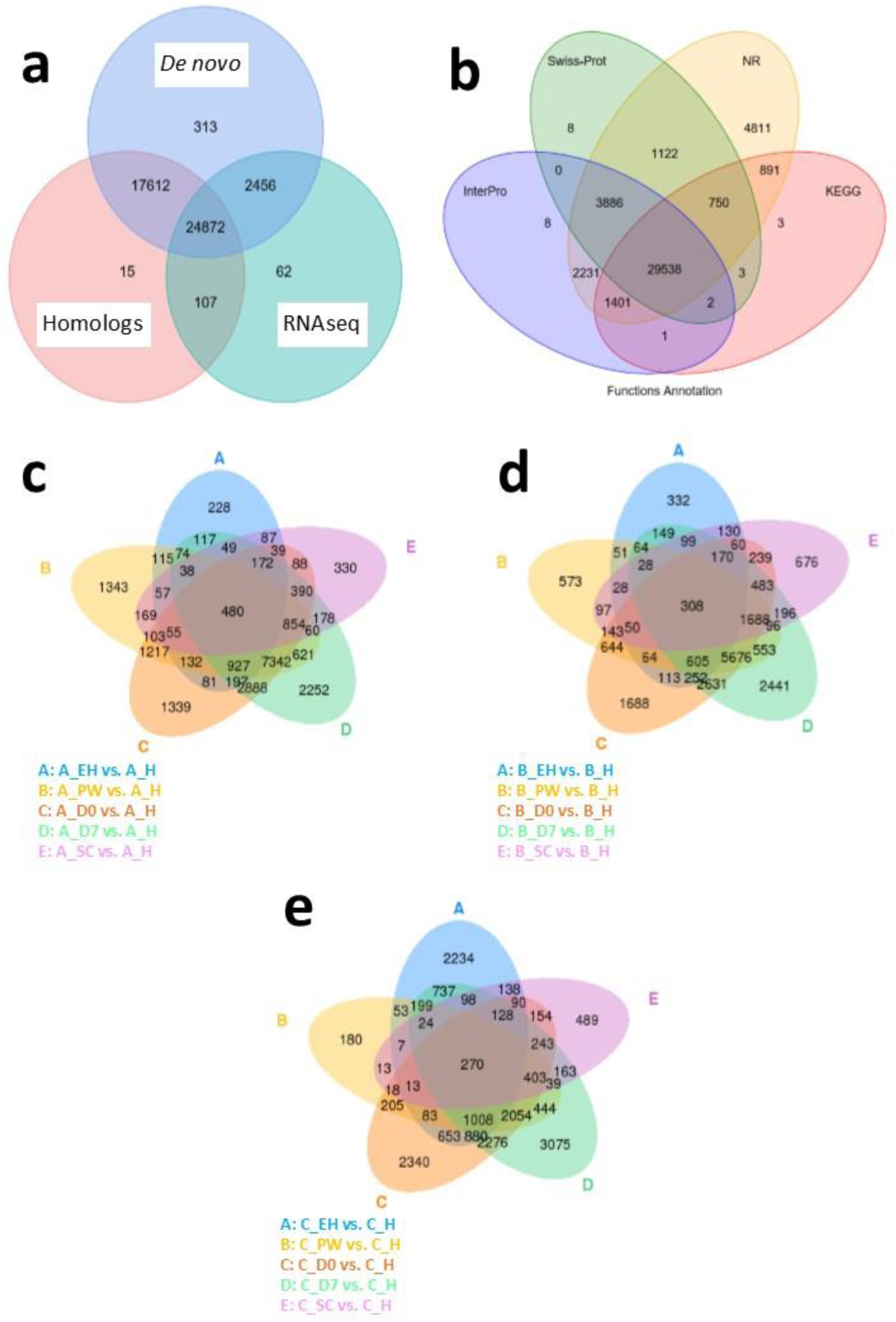
*Eruca sativa* genome annotation and differentially expressed gene number Venn diagrams for each inbred line. Venn diagrams of the *Eruca sativa* reference genome annotation gene identification sources (**a**) and functional annotation databases used to assign putative gene identities (**b**). Also shown are Venn diagrams of global differentially expressed genes (DEGs) at an early harvest (**EH**), second harvest (**SC**), pre-wash (**PW**), post-wash (**D0**), and seven-day shelf life (**D7**) time points relative to a first harvest (**H**) time point of three elite breeding lines: **A** (**c**), **B** (**d**), and **C** (**e**). The numbers of DEGs identified under each condition are contained within the ellipses and their overlaps.

### RNAseq analysis of *E. sativa* plants

RNA sequencing and bioinformatics was conducted on 18 plants from three elite inbred lines designated **A**, **B**, and **C**; giving a total of 54 plant samples. Time points corresponded to three harvest times (‘early harvest’ at 22 days after sowing, **EH**; ‘harvest’ at 30 days after sowing, **H**; ‘second cut’, **SC;** leaves harvested from the same **H** plants 43 days after sowing), and three consecutive postharvest time points (harvested at 30 days after sowing and designated: ‘pre-wash’, **PW**; ‘day 0’ of shelf life, 1 day post wash, **D0**; and ‘day 7’ of shelf life, **D7**). See Supplementary Figure S1 for a schematic of the experimental design.

After sample QC and clean-up over 2.6 billion clean paired-end reads were produced, averaging ∼49 million reads per sample. Q20 (<1% error rate) averaged 96.3%, Q30 (<0.1% error rate) averaged 90.8%, and GC content ranged from 44.5% to 47.4%.

### Global differential gene expression

The total numbers of differentially expressed genes (DEGs) for line **A, B** and **C** are presented in Figure 2c, 2d and 2e, respectively. Few significant DEGs were observed between **EH** and **H** samples for lines **A** and **B** (<333; Figure 2c and 2d), whereas they were observed at a higher rate in **C** (2,234; Figure 2e). This indicates a high level of plasticity of **C** across growth stages.

This trend was reversed at **PW**, where 180 DEGs were observed compared to **H** in **C**, and 1,343 were observed in **A**. During shelf life (**D0** and **D7**) **C** expressed a greater number of DEGs compared to **H**, than **A** or **B** (2,340 at **D0** and 3,075 at **D7**, respectively). By contrast, DEGs at **SC** were much less variable between the three lines (330 – 676) indicating a greater degree of uniformity of expression. In terms of DEGs different from **H** across all sample points, **A, B** and **C** had 480, 308, and 270, respectively.

Figure 3 presents the numbers of DEGs at each sample point between cultivars, along with the degree of spatial variation and separation using Principal Component Analysis (PCA). The close clustering observed is indicative of the high degree of reproducibility of expression seen between each biological replicate for each line tested (Supplementary Figure S2). It also emphasizes the differences in global expression patterns at any given ontological growth stage or postharvest time point between lines **A, B** and **C**. DEGs ranged from 17,167 at **PW** (37.78% of total genes; Figure 3b) to 22,482 at **EH** (49.48% of total genes; Figure 3a).

**Figure 3.**
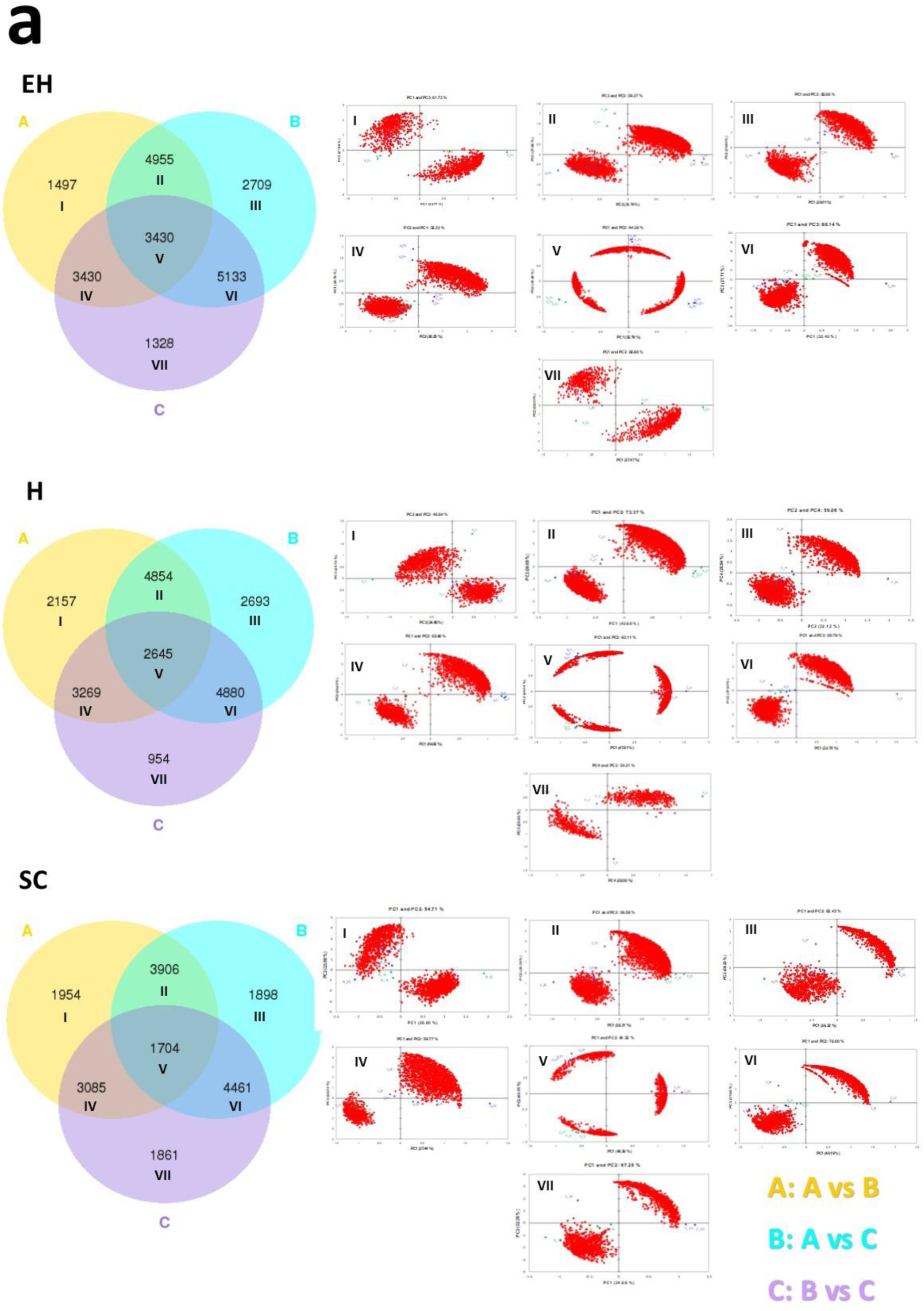
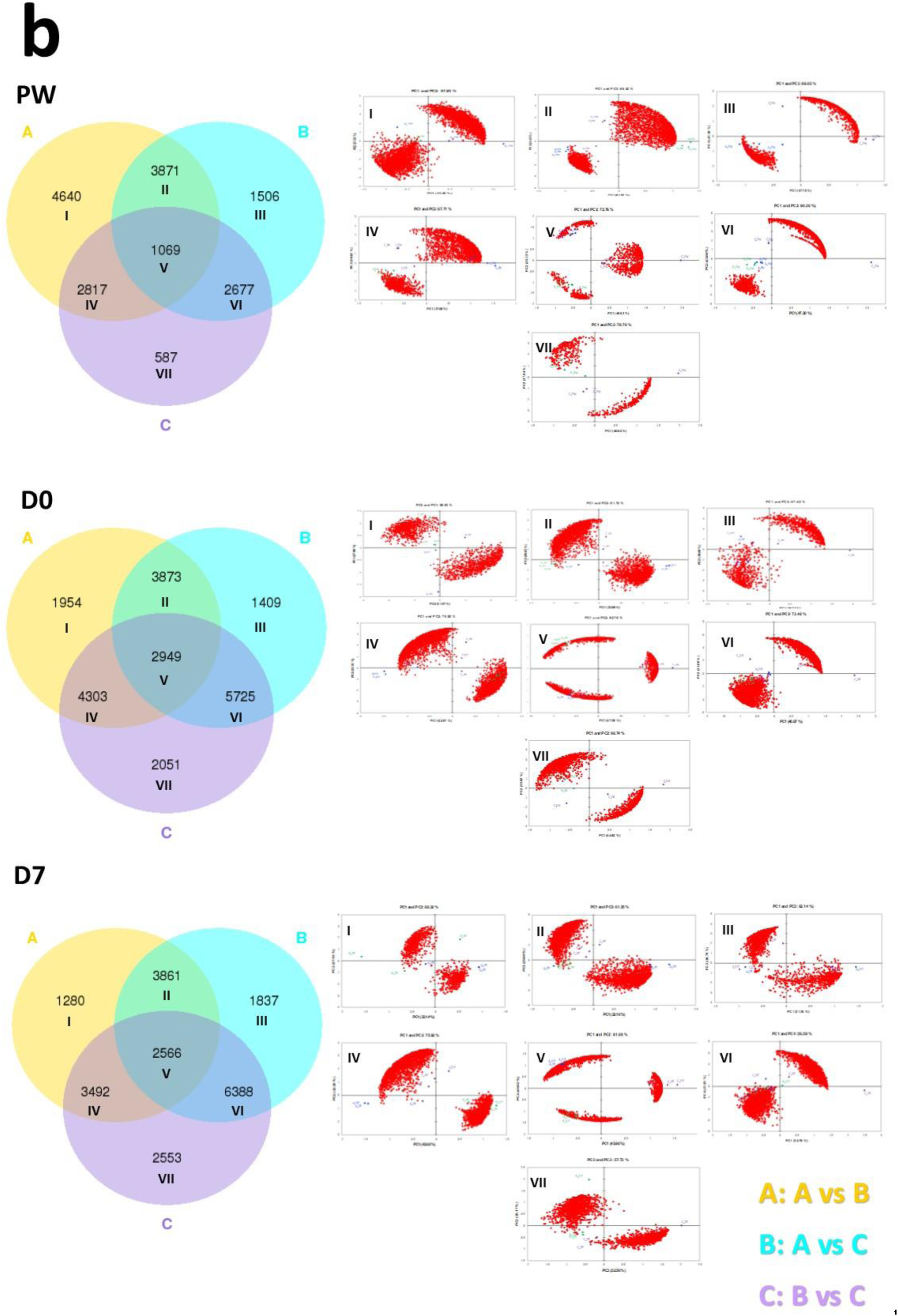
Venn diagrams of global differentially expressed genes between inbred lines with Principal Component Analyses. Venn diagrams of the numbers of differentially expressed genes (DEGs) found between three elite inbred lines of *Eruca sativa*. Different harvest ontogeny (**a**) and postharvest (**b**) time points are presented with accompanying Principal Component Analysis (PCA) plots that highlight the multidimensional separations of each DEG cluster between lines **A, B,** and **C**. Roman numerals are indicative of corresponding PCA analysis of each Venn diagram segment. Red data points within each loadings-scores biplot represent gene expression data (FPKM). Blue circles = **A**; green circles = **B**; purple circles = **C**. See insets for Venn diagram color coding. Abbreviations: early harvest (**EH**), harvest (**H**), second harvest (**SC**), pre-wash (**PW**), post-wash (**D0**), and seven-day shelf life (**D7**).

PCA of each of the DEGs (Figure 3) was used to scrutinize genes responsible for the highest degree of spatial separation according to factor scores. This analysis yielded 1,568 genes. Of particular note are several genes related to sulfur metabolism and GSL biosynthesis: *SLIM1a, MYB28b, GSTF11, IGMT4a, TGG1c, NIT2c, ESM1b, ESM1d, GTR2a, GTR2d, DHAR3, SiRb, SULTR2;2,* and *SULTR3;5a*.

### Sulfur and phytochemical composition of *E. sativa*

#### Sulfur content of *E. sativa*

Total sulfur content for each of the breeding lines is presented in Figure 4a. No significant differences were observed between lines and sample time points (*P* = 0.434). As will be discussed in the subsequent sections this observation is of significance.

**Figure 4.**
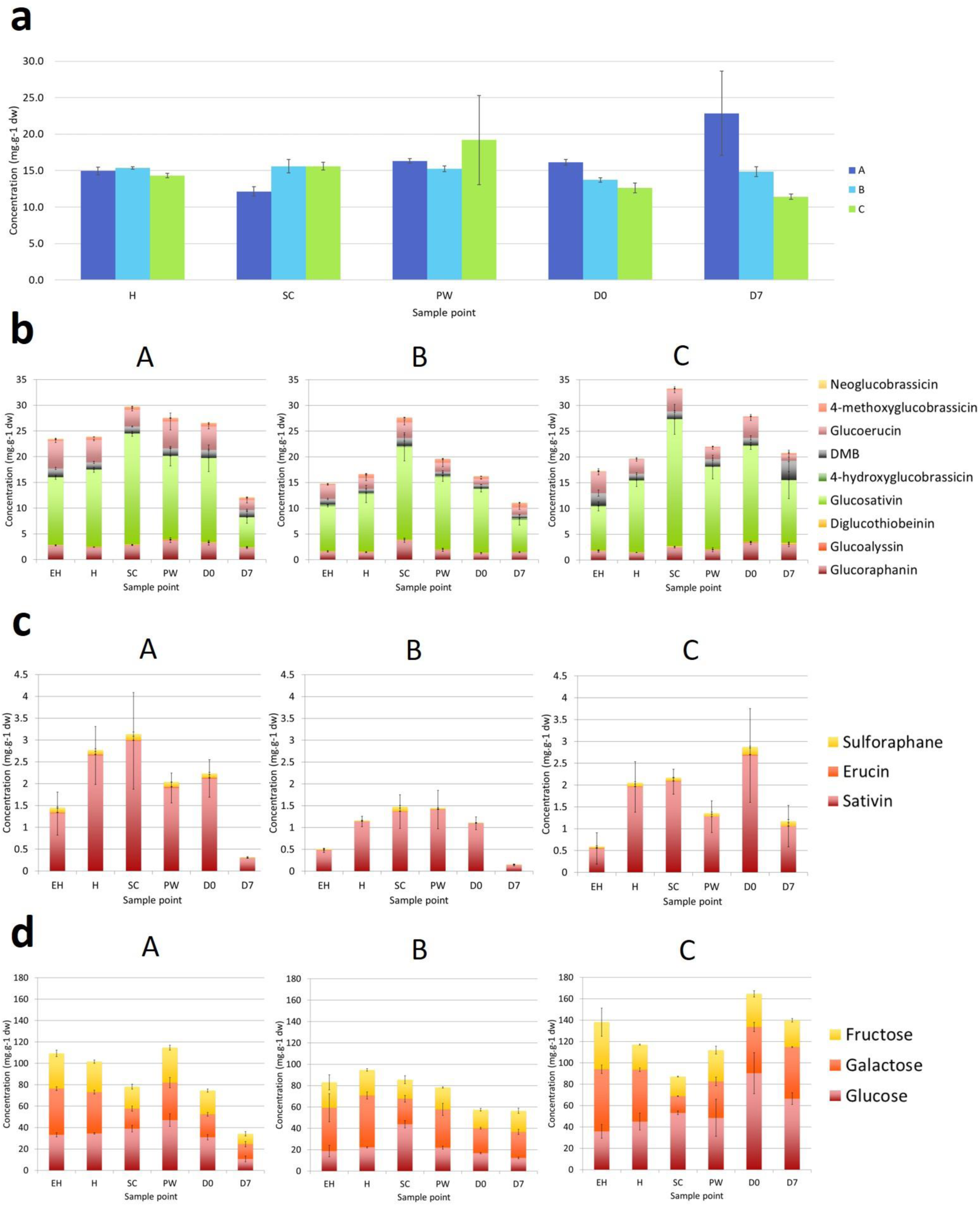
Elemental sulfur and phytochemical compositions of rocket inbred lines. Elemental sulfur (**a**), glucosinolate (**b**), glucosinolate hydrolysis product (**c**), and monosaccharide (**d**) concentrations observed in elite inbred lines of *Eruca sativa* (**A, B,** and **C**). Concentrations are expressed as mg.g^-1^ of dry weight. Error bars represent standard error of the mean of each analyte detected. See insets for compound color coding. For ANOVA and Tukey’s HSD pairwise significance values see Supplementary Data File S1. Abbreviations: early harvest (**EH**), harvest (**H**), second harvest (**SC**), pre-wash (**PW**), post-wash (**D0**), and seven-day shelf life (**D7**).

#### Glucosinolate profiles and contents of *E. sativa*

For each of the cultivars between the first (**H**) and second cuts (**SC**), an increase in total GSL concentrations was observed due to elevations of GSV (**A**, a 1.4-fold increase, *P*<0.0001; **B**, a 1.6-fold increase, *P*<0.0001; **C**, a 1.8-fold increase, *P*<0.0001) and GRA (**B**, a 2.6-fold increase, *P*<0.0001; **C**, a 1.8-fold increase, *P*<0.0001; Supplementary Data File S1). Line **C** produced the highest total concentrations of GSLs in **SC** (a 1.7-fold increase; *P*<0.0001), and line **B** also saw significant elevations compared to **H** (a 1.6-fold increase; *P*<0.0001).

Line **A** contained the greatest GSL concentrations compared to **B** and **C**, until **D7** where content declined significantly (a 0.6-fold decrease compared to **D0**, *P*<0.0001; Supplementary Data File S1). **C** by comparison contained high concentrations of GSLs during shelf life, peaking at **D0**, with a non-significant decrease at **D7** (0.3-fold reduction). This line did not demonstrate the same decline in GSLs towards the end of shelf life as in the other two, and displays a propensity for maintaining GSLs for longer into the shelf life period. We hypothesised that this was due to fundamental differences in expression of GSL biosynthesis genes pre and postharvest.

#### Glucosinolate hydrolysis product profiles and contents of *E. sativa*

Glucosinolate hydrolysis product (GHP) concentrations are presented in Figure 4c (see Supplementary Data File S1 for ANOVA and Tukey’s HSD significances). As with previous studies of rocket ^17^, three main GHPs were detected: sativin (a 1,3-thiazepane-2-thione; hydrolysis product of GSV; SAT), erucin (ITC of glucoerucin; GER), and SF. The fluctuations in total GHP concentration mirror those observed for GSLs, however the increases between **H** and **SC** are much less pronounced, with no significant differences between cuts.

As with GSLs, line **B** displayed the lowest concentrations of GHPs, whereas the differences between lines **A** and **C** are less apparent. The trend of reduction of GHPs over shelf life is also visible for lines **A** and **B**, though only significant in **B** (a 0.9-fold reduction, *P*<0.0001). Concentrations remained higher in line **C** (1.2 mg.g^-1^ dw, a 0.6-fold reduction from **D0**).

#### Monosaccharide profiles and contents of *E. sativa*

Monosaccharides are important in terms of sensory attributes and the masking of bitter and pungent sensory attributes in rocket ^5^ altering consumer perception and preference. Glucose is also known to influence stress responses and interact with MYB transcription factors ^18^ (Figure 1b). The concentrations of sugars observed in *E. sativa* lines are presented in Figure 4d (see Supplementary Data File S1 for ANOVA and Tukey’s HSD significances).

Unlike previous reports ^15^ the changes in sugar concentrations in this study were dynamic across each of the respective time points. Both lines **A** and **B** contained low total concentrations compared to line **C**. Line **B** displayed consistent concentrations, with no significant differences observed. **A** showed a similar trend to GSL and GHP concentrations by declining at the end of shelf life (**D7**; a 0.5-fold decrease from **D0**, *P*<0.0001).

Line **C** is distinct from the others in terms of its sugar profile and the relative differences between sample points. Concentrations increased postharvest (**D0** and **D7**; a 1.4 and 1.2-fold increase relative to **H**, respectively), perhaps owing to a breakdown of stored carbohydrate to facilitate respiration. Line **C** sugar content consists primarily of glucose, whereas **B** tended to have greater concentrations of galactose, and **A** was composed of similar amounts of each monosaccharide.

### Sulfur assimilation and glucosinolate biosynthesis pathway gene expression analysis

#### Sulfate assimilation gene expression

Figure 5a presents differential gene expression within the sulfate assimilation pathway of *E. sativa*. All significances quoted hereafter were at the *P*<0.001 significance level. In the primary stages of sulfur metabolism, assimilated sulfate is activated via adenylation to adenosine-5’-phosphosulfate (APS), catalyzed by ATP sulfurylase (ATPS) ^19^. In *E. sativa* four ATPS-encoding genes were identified: *APS1a, APS1b, APS2,* and *APS3* (Figure 1a). Very few significant DEGs were observed between sample points for each respective rocket line (see Supplementary Data File S2 for full values and statistics of each sample comparison). However, between **H**, **SC**, and **PW**, each respective line did show significant differential expression of ATPS genes.

**Figure 5.**
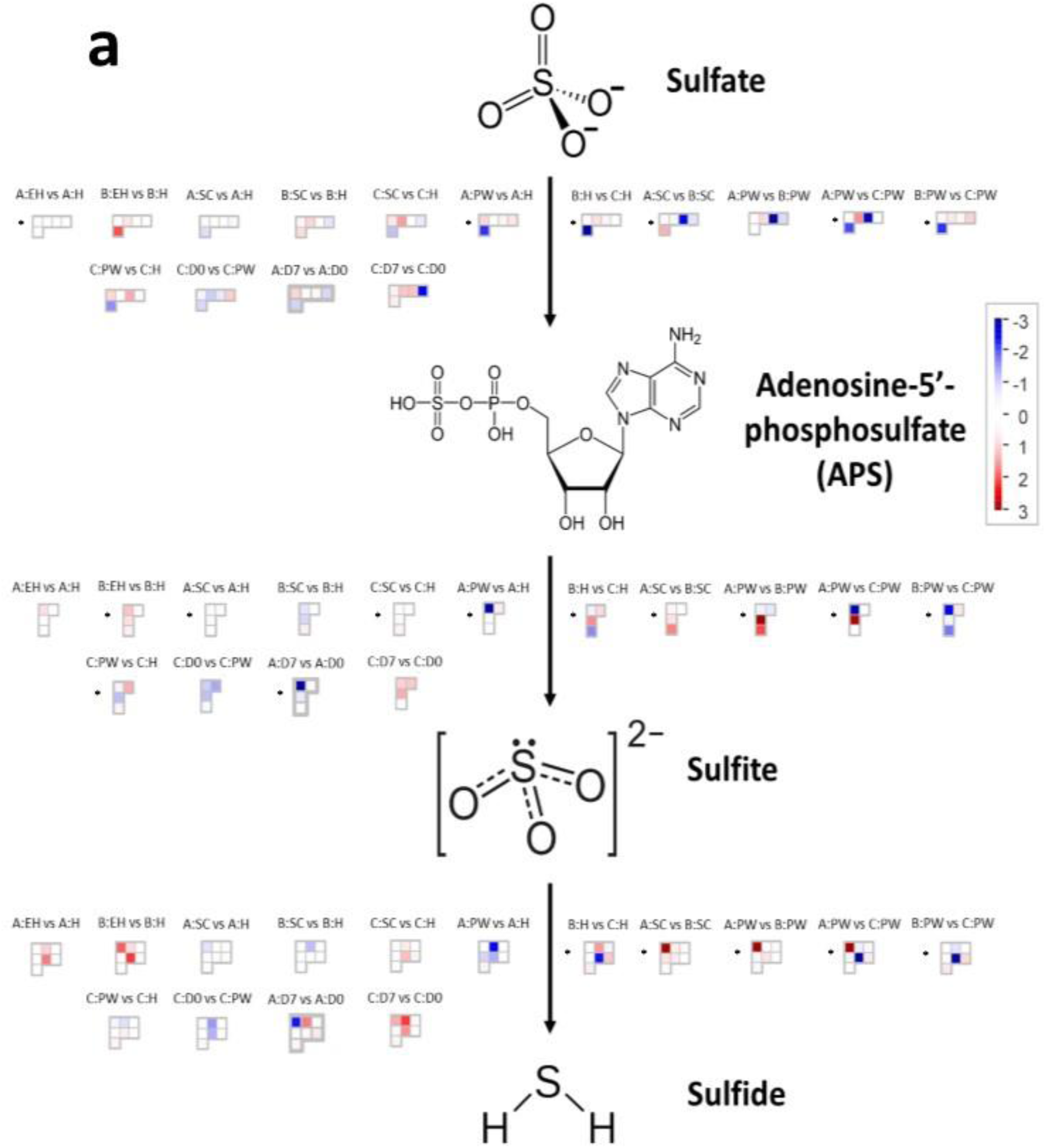
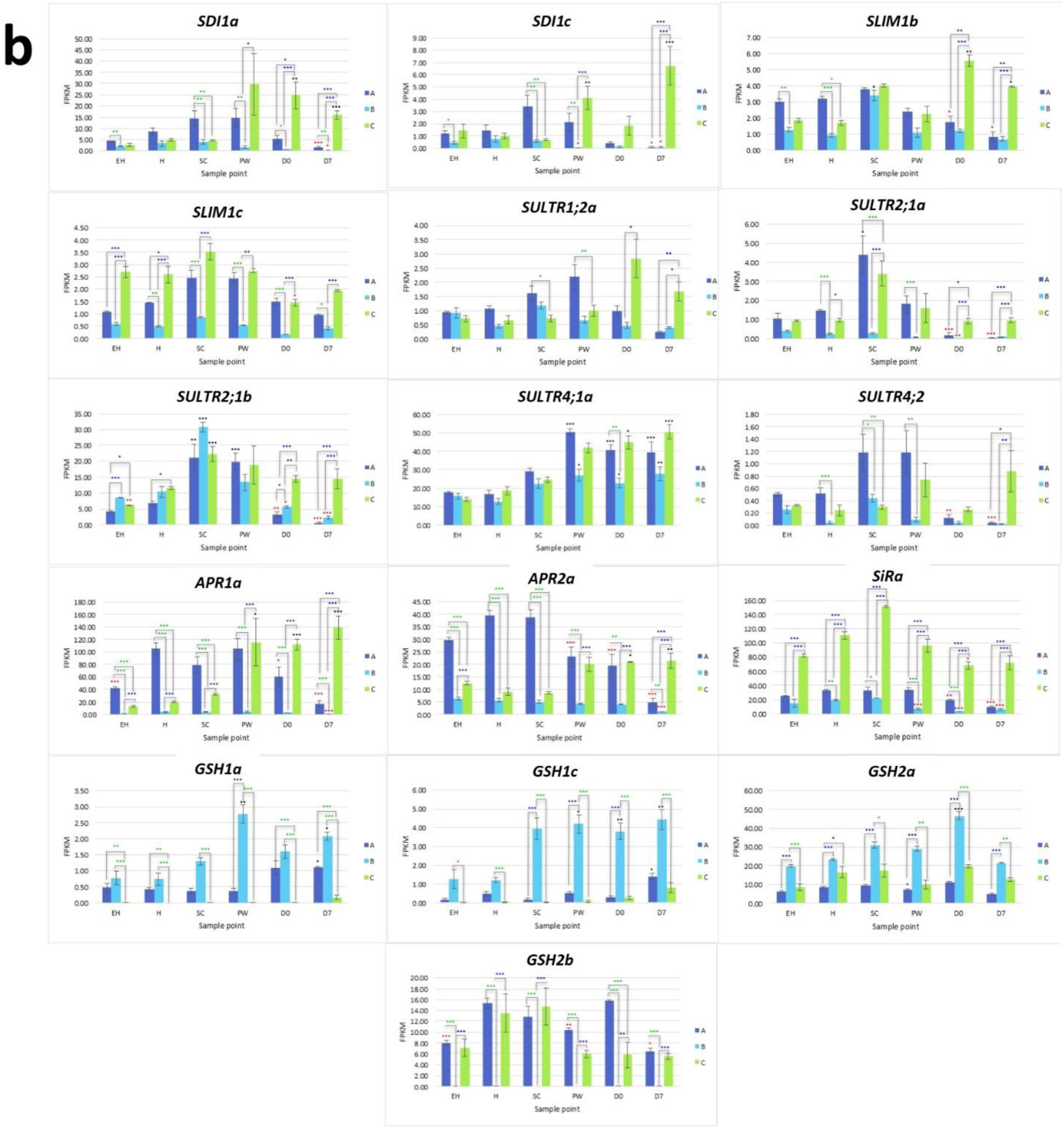
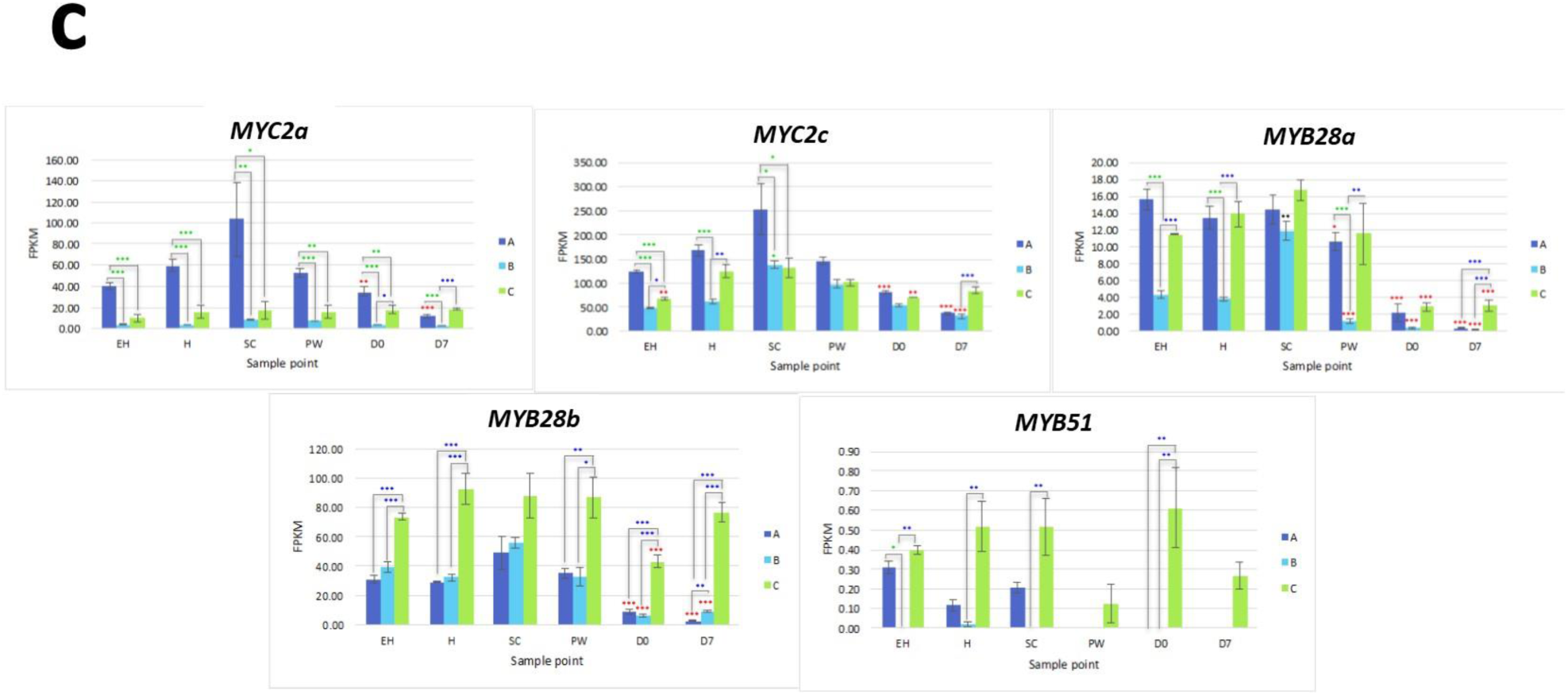
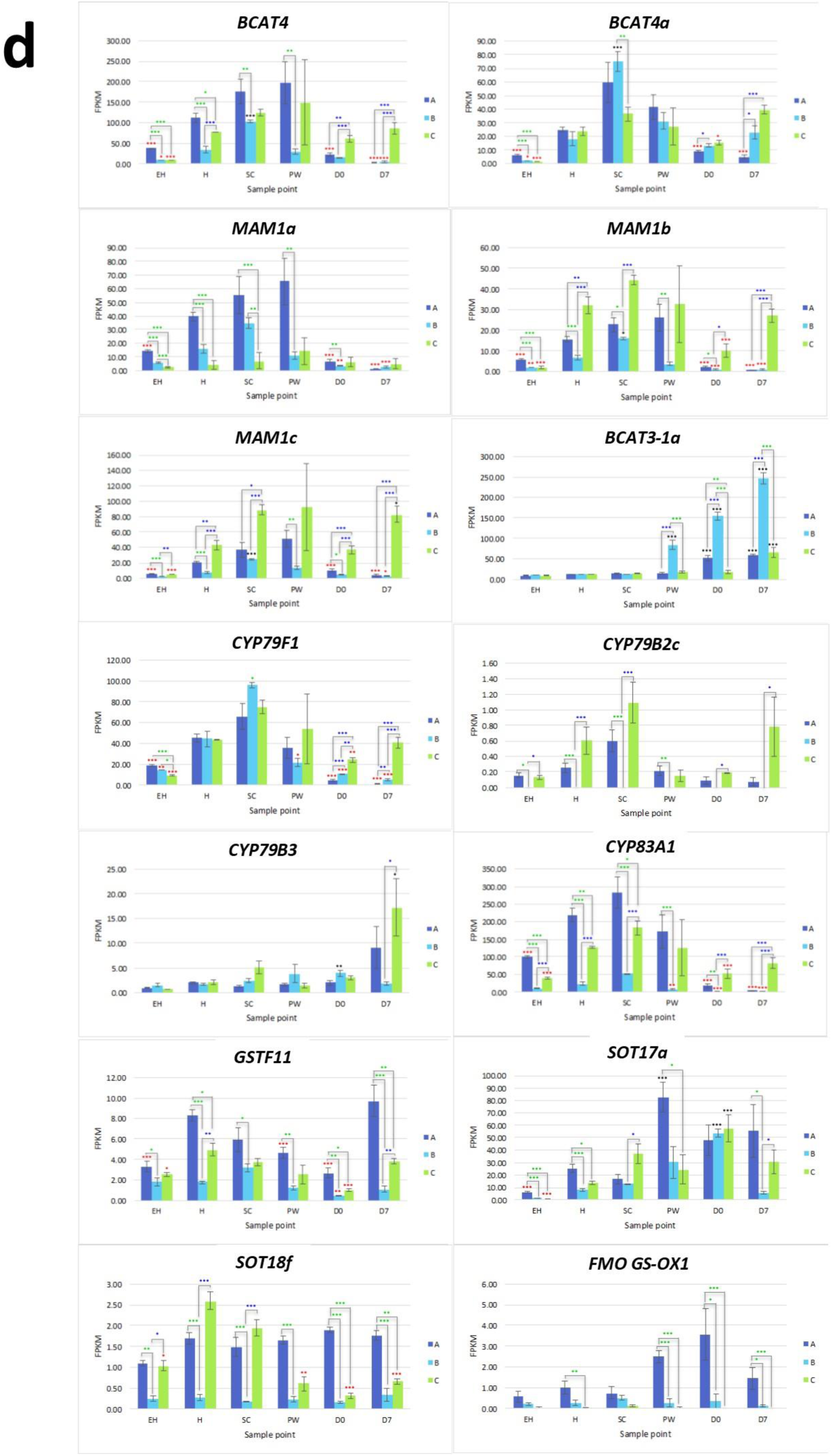
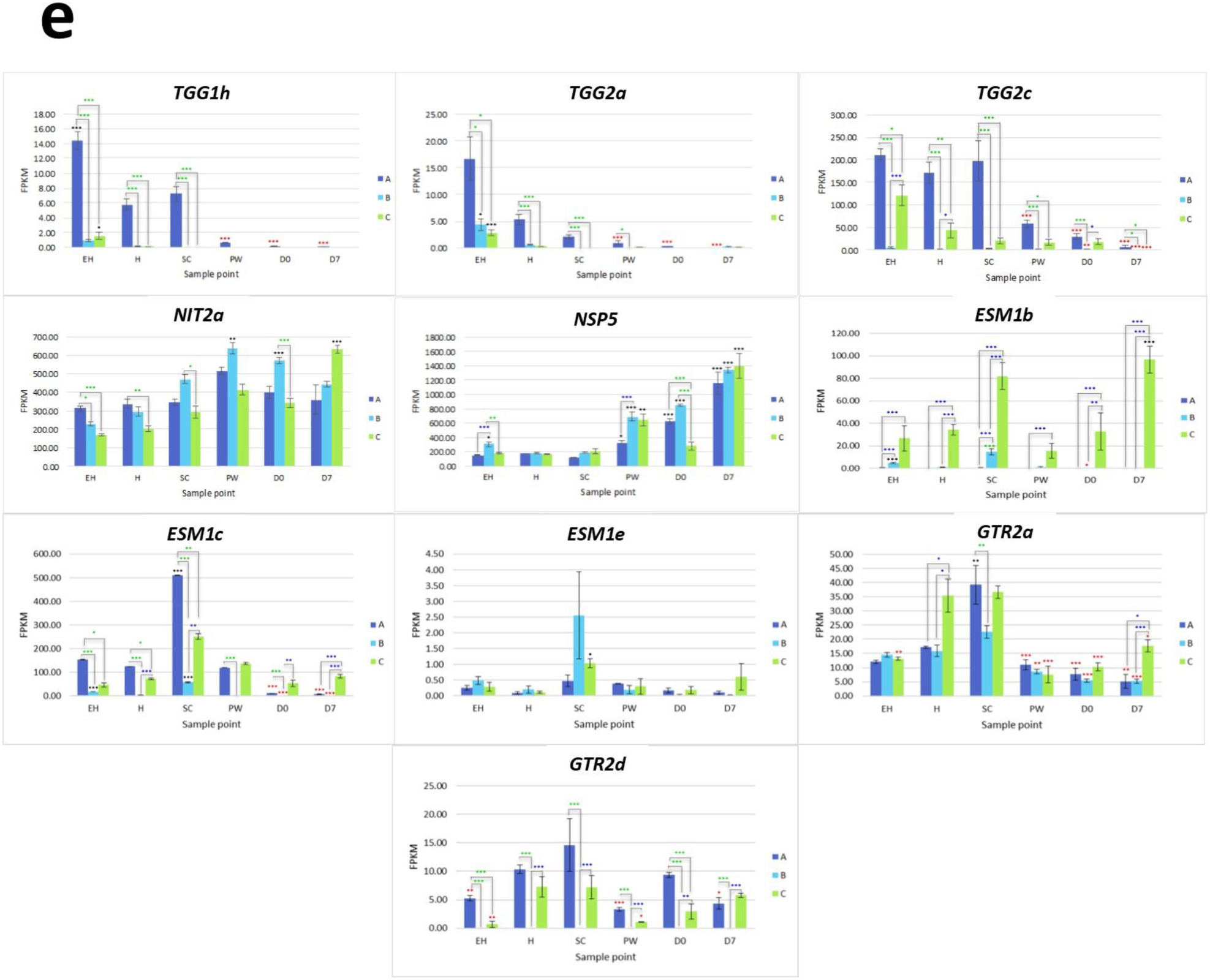
RNAseq expression data for sulfur metabolism and glucosinolate biosynthesis-related genes. RNAseq expression data (FPKM) for genes involved with sulfate assimilation (**a**), sulfate transport and redox response (**b**), glucosinolate transcription factors (**c**), glucosinolate biosynthesis (**d**), and glucosinolate hydrolysis and transport (**e**) in three elite inbred lines of *Eruca sativa* (**A** = dark blue; **B** = light blue; **C** = green). For (**a**) a custom MapMan (version 3.6.0RC1) annotation file of the *E. sativa* reference genome was created using Mercartor4 (version 1.0, plaBi dataBase, Institute of Biology, Aachen, Germany), and used to visualize differential expression of genes within the sulfate assimilation pathway. In (**b**) to (**e**), standard errors of the mean expression values are represented by error bars. Asterisks denote levels of significance of up and down regulation within sample points (between each inbred line) and relative to the point of harvest for each respective sample point: * = *P≤*0.05; ** = *P≤*0.01; *** = *P≤*0.001; green = significant up regulation between lines **A, B,** and **C**; blue = significant down regulation between lines **A, B,** and **C**; black = significant up regulation relative to **H**; red = significant down regulation relative to **H**. Abbreviations: early harvest (**EH**), harvest (**H**), second harvest (**SC**), pre-wash (**PW**), post-wash (**D0**), and seven-day shelf life (**D7**).

In the second stage of the pathway, APS is reduced to sulfite by adenosine-5’-phosphosulfate reductase (APR) genes ^20^. Four APRs were identified (*APR1a, APR1b, APR2a,* and *APR2b*) as well as six ARP-like genes (*APRL4, APRL5a, APRL5b, APRL5c, APRL7a,* and *APRL7b*). *APR1a* and *APR2a* showed significant differential expression across multiple samples and time points (Figure 4b). Line **B** displayed low relative expression of these genes compared to **A** and **C**. Line **C** exhibits significantly higher expression postharvest compared to **H**; 2.2 log2-fold (**D0**) and 2.7 log2-fold (**D7**) increases of *ARP1a*, and 0.9 log2-fold (**D0**) and 1.1 log2-fold (**D7**) increases of *APR2a* were observed. We hypothesise that this may be indicative of a greater ability to remobilize sulfate via APS genes to facilitate and maintain secondary metabolite biosynthesis for longer into shelf life.

Two copies of genes encoding sulfite reductase (SIR; *SiRa* and *SiRb*) were identified. *SiRa* showed significantly higher levels of expression in line **C** (Figure 5b). Line **C** had no significant change in activity of this gene relative to time point **H**, however both lines **A** and **B** had significantly lower expression postharvest (Figure 5b).

### Sulfur metabolism transcription, regulation, and transport gene expression

Three copies of *SDI1* (*SDI1a, SDI1b,* and *SDI1c*) and three copies of *SLIM1* (*SULFUR LIMITATION 1*, aka *ETHYLENE INDUCED 3*; *SLIM1a, SLIM1b*, and *SLIM1c*) were identified within the genome annotation. These genes are thought to play critical roles in the management and use-efficiency of sulfur in plants, and have been linked with optimization of GSL biosynthesis under S-limited conditions in *A. thaliana* ^10^.

*SDI1a* and *SDI1c* were differentially expressed between each line (Figure 5b, Supplementary Data File S2), with **C** having the highest levels of expression postharvest. It might be expected that each line would see a similar trend of expression over the course of shelf life, as additional sulfur is not obtainable; however only line **C** displayed this (Figure 5b). As shown in Figure 4a, sulfur content was not significantly different between any experimental stages or between breeding lines. Expression of MYB28 orthologs were not negatively associated with expression of *SDI1* gene copies. Previous research has shown that the SDI1 protein binds to MYB28, inactivating expression and reducing GSL biosynthesis ^21^. In *E. sativa* the opposite appears to be true, with significant positive correlations between respective expression of two MYB28 copies and *MYB29* with SDI1 copies (*SDI1a* and *MYB29*, *r* = 0.72; *SDI1b* and *MYB28a*, *r* = 0.507; *SDI1c* and *MYB28c*, *r* = 0.459; Figure 6). At **D7**, both **A** and **B** had significantly lower expression levels compared with **H** (a 2.2 and 3.7 log2-fold reduction of *SDI1a*, respectively; and a 3.9 and 3.2 log2-fold significant reduction of *SDI1c*, respectively).

**Figure 6.**
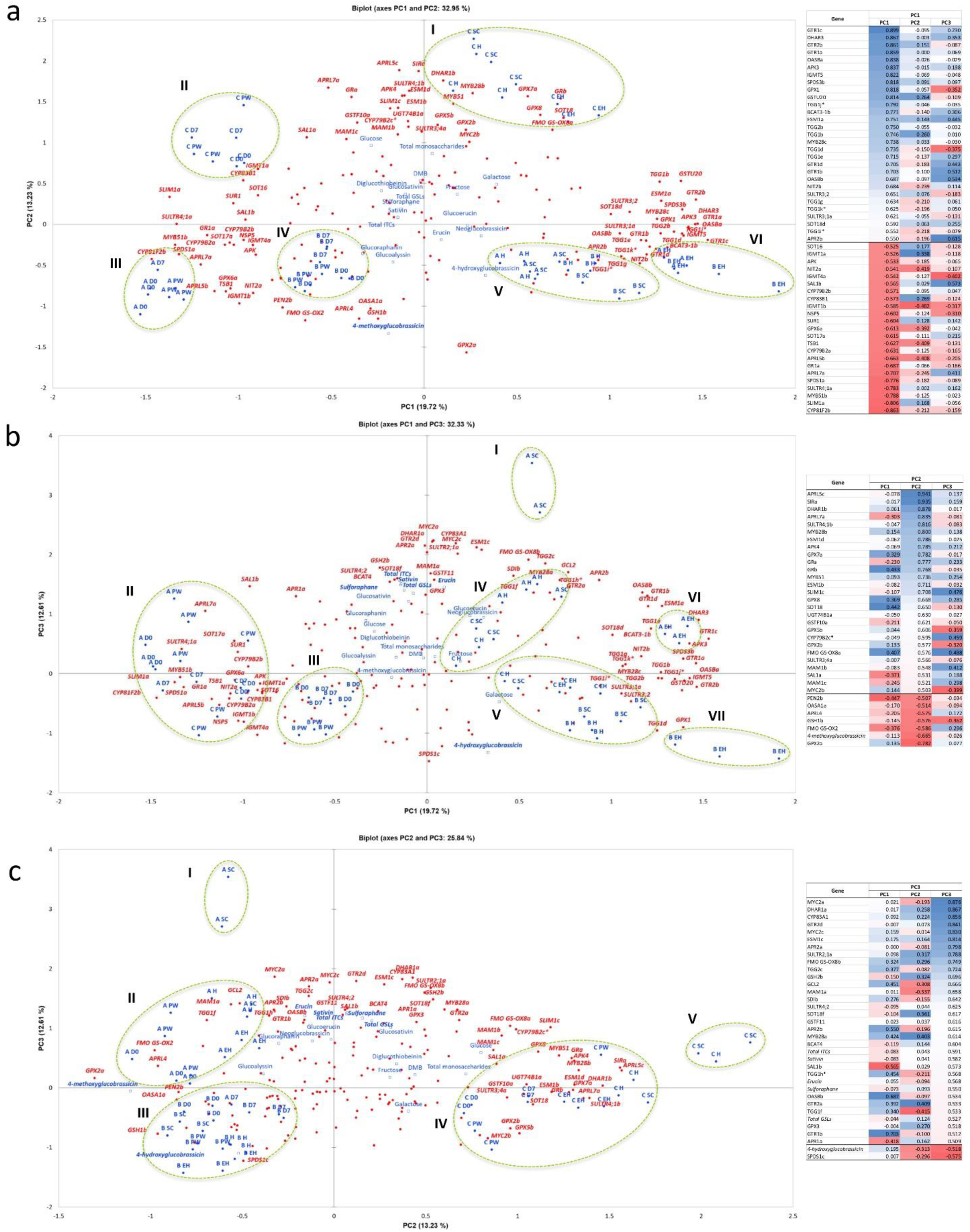
Principal Component Analysis of sulfur assimilation pathway, glucosinolate biosynthesis, and glucosinolate hydrolysis gene expression data. Principal Component Analysis of sulfur assimilation pathway, glucosinolate biosynthesis, and glucosinolate hydrolysis gene expression data (FPKM) for three *Eruca sativa* elite inbred lines (**A, B,** and **C**) across ontogenic and postharvest sample points. Biplot (**a**) displays Principal Components (PCs) 1 and 2, which represent 33% of variation within the data. Biplot (**b**) displays PC1 and PC3, explaining 32.3% of variation within the data; and (**c**) displays PC2 and PC3, explaining 25.8% of the variability. The PCA plots presented are the results of Varimax rotation. Each biplot is accompanied by a factor loadings table sorted according to PC1 (**a**), PC2 (**b**), and PC3 (**c**); italics denotes a supplementary variable, and * = a putative novel gene within the reference annotation. Blue coloration denotes high factor loading scores, red denotes low. Only genes with loading values >0.5 were included, and each is represented within the biplots in red (bold italics). Red circles represent individual genes included in the analysis (*n* = 177). Blue circles represent sample point variables and have accompanying labels (blue bold). Blue squares denote phytochemical data regressed onto the PCA as supplementary variables. Bold data labels indicate phytochemical components with >0.5 factor loadings scores. Green dotted ellipses denote clusters of variables and are numbered using Roman numerals, which are quoted within the text. Abbreviations: early harvest (**EH**), harvest (**H**), second harvest (**SC**), pre-wash (**PW**), post-wash (**D0**), and seven-day shelf life (**D7**).

A similar pattern was observed for *SLIM1b*, where expression in **C** was significantly higher at time points **D0** and **D7** relative to **H** (a 1.4 and 1.1 log2-fold significant increase, respectively). Expression of *SLIM1c* by comparison was not significantly different for each respective plant line between time points, but there were clear and significant differences in expression between lines (Supplementary Data File S2). Line **C** had highest expression of this gene, followed by **A**; with **B** having significantly lower expression (Figure 5b). Previous studies have shown that SLIM1 down regulates APK gene expression and GSL biosynthesis as a way of conserving sulfur for primary metabolism ^21^. Our data suggest that this is only the case between *SLIM1a* and *APK3* (*r* = −0.597, *P*<0.001; Figure 6). *SLIM1a* expression was positively (and significantly) correlated with *APK* expression (*r* = 0.521), and *SLIM1b* and *SLIM1c* with *APK4* (*r* = 0.575 and 0.698, respectively; Figure 6). This suggests *E. sativa* has a complex and interacting network of sulfur metabolism genes, where functions may not be analogous to those found in *A. thaliana*.

16 sulfur transport (*SULTR*) genes were identified; of note were *SULTR1;2a, SULTR2;1a, SULTR2;1b, SULTR4;1a,* and *SULTR4;2*. *SULTR1;2a* has been associated with the uptake of environmental sulfate in root tissues (Supplementary Data File S2), but low levels of expression were detected in leaf tissues. Postharvest, line **C** had differential expression of this gene compared to **A** and **B** in **D7** samples (Figure 4b). This was more pronounced for *SULTR2;1a* and *SULTR2;1b*, and both **A** and **B** had significant reductions in expression at **D0** and **D7** relative to **H**. **SC** samples showed significant increases relative to **H**, with the exception of *SULTR2;1a* in **B**.

*SULTR4;1* and *SULTR4;2* genes also had distinct patterns of expression between lines. *SULTR4;1a* saw significant increases in expression in postharvest samples relative to **H** (Figure 5b). Line **A** had higher expression of *SULTR4;2* during growth before declining significantly post-wash (**D0**). The opposite trend was seen in **C**, where gene expression peaked at **D7**. These data are suggestive of more active intra-leaf sulfur transport in line **C** postharvest, and may be associated with the higher expression of *APR, SIR, SDI1* and *SLIM1* genes to facilitate more efficient S utilisation during this period.

### Glutathione synthesis

With the exception of *GSH2b*, glutathione synthase genes were most highly expressed in rocket line **B**, with significant increases observed postharvest (Figure 5b). Line **A** and **C** were unchanged between sample points for these genes, but had a marked difference in expression for *GSH2b* relative to each other. **B** had negligible levels of *GSH2b* expression.

As both glutathione and secondary S-containing metabolites, such as GSLs, have been associated with antioxidant responses ^21^ the differences observed between each of the lines in terms of both GSL concentrations and glutathione-related gene expression, may be indicative of different adaptive metabolic strategies for dealing with oxidative stress postharvest. Lines **A** and **C** favor secondary sulfur metabolism and the synthesis of GSLs, and **B** favors primary sulfur metabolism and glutathione synthesis.

### Glucosinolate-related transcription factors

*MYC2a* and *MYC2c* were highly expressed in line **A**, and had uniform patterns of relative expression. **SC** had the highest expression values for this line, suggesting a general response to mechanical wounding and stress, however this was not significantly different from **H**. The only significant difference for *MYC2c* between **H** and **SC** was in line **B** (a 0.7 log2-fold increase; Figure 5c).

*MYB28a* and *MYB28b* display high degrees of differential expression between each rocket line. While **A** has high expression of *MYB28a* in samples **EH, H, SC**, and **PW**, it is has by comparison lower expression of *MYB28b* compared to **C** (Figure 5c). **C** on the other hand has relatively high expression for both of these TFs, and displays significantly higher expression postharvest, up to and including **D7**. Combined with what is known about theses TFs in other Brassicaceae species, it is likely that the differences in GSL concentrations observed postharvest are linked to the differential expression of *MYB28a* and *MYB28b* between the respective lines.

### Glucosinolate biosynthesis

Rocket contains two genes encoding BCAT4, and two genes encoding BCAT3; converting 2-oxo acids to homomethionine and dihomomethionine. *BCAT3-1a* displayed no significant variation between lines during growth, but saw significant increases for all (compared to **H**) at **D0** and **D7** (Supplementary Data File S2). The most marked and significant increase was in **B**. It is unclear how this ‘preference’ for BCAT3 activity over BCAT4 is regulated or affects the synthesis pathway, but the relative and respective activity of these genes is correlated with GSL content.

Only orthologs of *MAM1* were identified, and each of the three copies had differing expression patterns (Figure 5d). Line **A** displayed higher relative expression of *MAM1a*, whereas **C** had greater expression for *MAM1b* and *MAM1c*. **B** however maintained low expression for all three of these genes. **A** had reduced expression activity during shelf life, whereas in **C**, levels were significantly higher compared to **H** (Figure 5d).

One *CYP79F1* homolog was found in rocket, with no expression found for a corresponding *CYP79F2* gene. The lack of a *CYP79F2* homolog in rocket may be suggestive of a loss of function, and/or redundancy with other enzymes. Of note for *CYP79F1* expression was the significant differences observed between **EH** and **H**, indicating that earlier harvests of rocket leaves may have a reduced ability for GSL biosynthesis compared with later ones and second cuts (**SC**). Expression was significantly greater in **C** during shelf life. In the conversion of aldoximes to nitrile oxides, *CYP83A1* expression was higher in **A** and **C** than **B**, with line **C** having significantly higher expression in shelf life samples (Figure 5d).

### Glucosinolate hydrolysis

11 *TGG1* (myrosinase) orthologs, and three *TGG2* (Supplementary Data File S2) were identified within the annotation. Some of these genes appear to have differential expression according to ontogeny and shelf life, with some copies expressed at **EH** with none during postharvest (e.g. *TGG1h, TGG2a,* and *TGG2c*; Figure 5e). Others however display the inverse of this, with increased relative expression postharvest (*TGG1a*; Supplementary Data File S2). It is known that myrosinases are functionally redundant, however it has also been noted that their activity and specificity is linked with developmental processes, and may explain some of the high levels of expression observed at **EH**.

An explanation for the lack of nitrile GHPs in rocket may be that the high expression of *NSP5* is inhibited by the five *EPITHIOSPECIFIER MODIFIER* 1 (*ESM1*) orthologs found in the rocket genome. These proteins are known to inhibit the action of NSPs and promote ITC formation. Expression was significantly greater in line **C** for *ESM1b* (Figure 5e) at all sample points, and fits with the observed pattern of sustained GHP formation postharvest. The lower activities in **A** and **B** did not however correspond to a reciprocal decrease in the relative concentrations of GHPs, and neither were nitrile concentrations at anything above trace levels. Much further work is needed to explain the genetic regulation of GHP formation in rocket and the high expression of *NSP5*.

### Glucosinolate transporters

Eight GSL transporter genes were identified in the rocket annotation; four *GTR1* and four *GTR2* homologs. These genes are involved in leaf distribution and long-range phloem GSL distribution, respectively. Expression of *GTR2a* and *GTR2d* were significantly correlated with GSL abundance and GHP formation in the analysed leaf tissues. Of particular note is that **B** had no expression of *GTR2d* at any of the sample points, indicating that the gene may be non-functional, and contribute to the overall low concentrations of GSLs observed. The exact site(s) of GSL synthesis has not been established in *E. sativa*, however it is known that transport from the root tissues to leaves occurs in other Brassicaceae. If this transport system is impaired in **B**, this would explain the significantly lower abundance of GSLs observed in leaves (Figure 4b). Coupled with the high expression of glutathione-related genes and similar sulfur content of **B** compared to lines **A** and **C**, the inactivity of this gene copy may have significant effects on leaf sulfur transport, metabolism, and antioxidant response. The lower GSL content in leaves may therefore be compensated by increased glutathione synthesis.

### Principal component analysis of sulfur and glucosinolate metabolism genes

Hereafter, only correlations significant at the *P*<0.001 level are presented and discussed. *SULTR4;1a* was significantly correlated with GRA concentrations (*r* = 0.577), which is associated with shelf life samples for lines **A** and **C** (Figure 6b, cluster **II**). Figure 6a and 6b show a distinct separation between ontogenic and shelf life samples along PC1. The increased expression of sulfur transport genes such as this postharvest may provide some explanation as to why GSL concentrations increase in the initial stages shelf life (**PW**), as S may be re-mobilized to facilitate biosynthesis. Efficient transport and storage of sulfur pre-harvest may also facilitate better retention and decreased degradation of GSLs postharvest. This can be seen in Figure 6b (**V**) where *SULTR3;1a* and *SULTR3;2* are associated with pre-harvest expression.

SF and SAT concentrations were significantly correlated with *APR2a* gene expression (Figure 6c **I** and **II**; *r* = 0.58, SF; *r* = 0.464, SAT) and associated in particular with **A** ontogenic samples and **PW**. *APR2* is known to contribute to sulfur accumulation and homeostasis, as well as facilitating cysteine synthesis, and is associated with increased myrosinase activity and GSL recycling. Line **A** (on average) contained the highest ontogenic concentrations of GRA, SF, GSV, and SAT (Figure 4b and 4c); this is supported by a significant correlation and association with GSL-related transcription factors *MYB28a* (*r* = 0.486, SF; *r* = 0.53, SAT), *MYC2a* (*r* = 0.596, SF; *r* = 0.626, SAT) and *MYC2c* (*r* = 0.584, SF; *r* = 0.583, GSV; *r* = 0.634, SAT; Figure 6c **I** and **II**), as well as a drought tolerance-related gene *SAL1b* (*r* = 0.595, GRA; *r* = 0.547, SF; *r* = 0.499, SAT; Figure 6a **II, III** and **IV**, 6c **II**). **A** was also associated with increased activity of *MAM1a* (Figure 6c **II**), facilitating greater GRA biosynthesis through chain elongation. It may be that lines **A** and **C** have increased relative GRA concentrations at **EH** and **H**, but preferentially express *MYB28c* and *MYB28b*, respectively. It is unknown if the function of each MYB28 TF are redundant in rocket, but these data would suggest that there is some clear overlap of function, though the expression of *MYB28b* is associated with increased GSL biosynthesis postharvest (Figure 5c).

The lower relative expression in line **B** for many of these genes is consistent with its lower GSL and hydrolysis product concentrations, irrespective of sample point. GRA/SF, GSV/SAT, and GER concentrations were significantly and negatively correlated with *SPERMIDINE SYNTHASE 1c* (*SPDS1c*; *r* = −0.622, GRA; *r* = −0.614, SF; *r* = −0.6, GSV; *r* = −0.454, SAT; *r* = −0.604, GER), which had a high degree of co-separation in all **B** samples (Figure 6c **III**). This association may be related to increased primary S metabolism and reduced partitioning of methionine for secondary S metabolites (Figure 1a).

## Discussion

### *E. sativa* has a distinct and complex glucosinolate pathway

Gene orthologs have undergone numerous duplications in *Eruca*, and it is not clear what the function(s) of these numerous copies may be. It may be the result of segmental duplications within the genome, such as those observed in the *Brassica* A genome ^22^, and future, more detailed studies of the *Eruca* genome structure may reveal the nature and number of any such events. Genes such as SOTs, FMO GS-OXs, myrosinases (TGGs), *ESM1*s, and GSL transporters all have several copies, and it has yet to be determined if these perform the same function as in *Arabidopsis,* or have evolved new ones that are responsible for the unique GSL profile of rocket. Examples of this include increases in copy numbers compared to *B. rapa* and *B. oleracea*. In these two species two copies of *SOT18* have been identified ^23^, whereas *E. sativa* has seven. *B. rapa* has two copies of FMO GS-OX genes, and salad rocket has at least ten. The related *Diplotaxis tenuifolia* (“wild” rocket) transcriptome has been reported to contain three copies of *MYB28* ^24^, and is consistent with the hypothesis that duplication occurred after their divergence with a common ancestor within the *B. oleracea* lineage.

Both *Arabidopsis* and *B. rapa* have four myrosinase gene copies, and *B. oleracea* has six ^23^. Our data indicate that *Eruca* has at least 14 copies, as well as two copies encoding PEN2 myrosinase. There has evidently been a massive diversification and duplication of these enzymes in rocket, but it has yet to be established if this is reflected in functionality and spatial expression.

### Glutathione metabolism competes with glucosinolate biosynthesis for sulfur

As shown in Figure 4a, the content of sulfur between the three tested breeding lines was not significantly different. In light of the observed differences in gene expression and GSL accumulations, we theorize that primary and secondary sulfur metabolism pathways ‘compete’ for assimilated environmental sulfur. As content was not significantly different postharvest (**PW, D0,** and **D7**) compared to pre-harvest first cut (**H**) in any of the breeding lines, the degree of remobilization and ability to synthesize/recycle GSLs is under strict genetic control. The exact reasons why each line differs in this respect are unclear, but as shown in Figure 4b, the amount of total sulfur assimilated during growth is not reflected in the postharvest concentrations of GSLs. Line **B** displays this: it contains significantly no more or less sulfur than lines **A** or **C**, yet synthesizes far fewer GSLs.

We hypothesise that the lack of expression of *GTR2d* impairs GSL transport from major sites of synthesis (possibly in the roots) thereby forcing leaf tissues to cope with oxidative stress via glutathione synthesis and preferential shunting of sulfur into the primary metabolism pathway. This is evidenced by the significantly higher expression of glutathione synthetase (*GSH2a*) and glutamate-cysteine ligase genes (*GSH1a* and *GSH1c*). Lines **A** and **C** by contrast may cope with oxidative stress postharvest by sustaining GSL biosynthesis. As **C** contained greater concentrations of monosaccharides, we theorize that this facilitates GSL metabolism for longer, reducing oxidative stress, and delaying the onset of senescence.

### Second cut rocket has greater uniformity of gene expression and glucosinolate content than first cut

Anecdotal evidence has suggested that the second cut (**SC**) is more consistent in terms of taste and flavour than the first (**H**). For the first time we present transcriptomic and phytochemical evidence to support this assertion. All cultivars saw increases in GSL abundance in the second cut, and the shift is best exemplified by line **B**. First cut (**H**) samples were low in expression of numerous GSL biosynthesis related genes that later saw significant increases at **SC**. This had the result of making its expression profile more similar to that of **A** and **C**, along with comparable concentrations of GSLs.

When the respective monosaccharide profiles are taken into account, it can be seen that there is a general decrease in sugars at **SC**. This then leads to a shift in the ratio between monosaccharides and GHPs, making the **SC** samples more pungent. Although flavour was not tested in this study, our previous work has highlighted the importance of GHP:sugar ratios in determining pungency ^25^.

## Materials and Methods

### Plant material for genome sequencing

Three elite inbred lines of salad rocket were produced through self-pollination for five generations at Elsoms Seeds Ltd. (Spalding, UK) from 2010-2016, giving an estimated inbreeding coefficient of 0.969 ^26^. Each line was derived from germplasm accessions obtained from the Leibniz-Institut für Pflanzengenetik und Kulturpflanzenforschung (IPK Gatersleben, Germany). For reasons of commercial sensitivity these lines (**A**, **B**, and **C**) and their lineage will not be identified.

For genome sequencing, plants of each line were grown under controlled growth room conditions, and leaf samples had DNA extracted and sent to the Earlham Institute for QC analysis. Samples were quantified using a Qubit fluorometer and dsDNA assay kit (ThermoFisher Scientific, Loughborough, UK) and assessed for quality using NanoDrop (ThermoFisher Scientific; according to 260/280 and 260/230 ratios).

### Genome sequence library preparation and assembly

DNA sequencing and assembly was performed as a service by the Earlham Institute (Norwich, UK). *De novo* genome sequencing and assembly was performed using PCR free paired-end (PE) and LMP sequencing. After DNA sample QC, line **C** was selected for sequencing and reference genome assembly. One PCR free PE library was constructed from gDNA, and sequenced on one lane of an Illumina HiSeq2500 in rapid run-mode (v2) using 250 bp PE reads. LMP sequencing was also conducted using one set of Nextera libraries (Illumina) from gDNA, and sequenced on one lane of an Illumina MiSeq with 250 bp PE reads. After data QC and assembly of the high coverage PE library, LMP libraries were mapped to determine their suitability for assembly improvement. Three additional libraries were selected and re-sequenced to a higher depth of coverage on a single lane of an Illumina HiSeq2500 in rapid run-mode, to again yield 250 bp PE reads.

### Genome sequencing bioinformatics

FASTQ files were converted to BAM format using PicardTools (v1.84, http://broadinstitute.github.io/picard/; FastqToSam option) and then assembled using DISCOVAR *de novo* sequence assembler (build revision 52488) ^27^. All LMP libraries were processed using NextClip ^28^ to analyse and create a high quality read subset for scaffolding the DISCOVAR-assembled sequences. SOAP ^29^ and SSPACE ^30^ were used to scaffold the DISCOVAR assembly using data from three of the NextClip-processed LMP read libraries.

### Genome annotation

Annotation was performed by Novogene Co. Ltd. (Hong Kong). A homology and *de novo*-based approach was taken in order to identify TEs. The homology-based approach used known repetitive sequence databases: RepBase ^31^, RepeatProteinMask, and RepeatMasker (http://www.repeatmasker.org/). *De novo* repeat libraries were created using LTR_FINDER ^32^, RepeatScout (http://www.repeatmasker.org/), and RepeatModeler (http://www.repeatmasker.org/RepeatModeler.html).

An integrated approach was taken to compute consensus gene structures, such as cDNA, proteins in related species, and *de novo* predictions. The homology-based approached used the related genomes of *A. lyrata, A. thaliana, B. napus, Boechera stricta, Capsella rubella,* and *R. sativus* (Supplementary Table S3) to compare against *E. sativa* to find homologous sequences, and predict gene structures (using BLAST and genewise) ^33, 34^. *Ab initio* statistical models were also used to predict genes and their intron-exon structures; e.g. Augustus ^35^, GlimmerHMM ^36^, and SNAP (http://homepage.mac.com/iankorf/). EVidenceModeler (EVM) ^37^ software was then used to combine *ab initio* predictions, protein and transcript alignments, and RNAseq data into weighted consensus gene structures. Lastly, PASA was used to update the consensus predictions by adding UTR annotations and models for alternative splicing isoforms. All predicted proteins were functionally annotated using alignments to SwissProt, TrEMBL ^38^, KEGG ^39^, and InterPro ^40^ (Figure 2b).

The full reference genome sequence and annotation can be found in the European Nucleotide Archive (Assembly accession no: GCA_902460325; Study ID: PRJEB34051; Sample ID: ERS3673677; Annotation accession number ERZ1066251).

### Plant material growth and collection for RNA, elemental, and phytochemical analyses

For RNAseq analyses seeds were sown in a random order in seedling compost, and raised under controlled environment conditions in plastic trays inside a Weiss-Technik Fitotron cabinet (Weiss-Technik UK Ltd., Loughborough, UK). Daytime temperature was set to 20 °C, and nighttime temperatures to 14 °C (long day cycle; 16 h light, 8 h dark). Light intensity was set at 200 μmol m^-2^ s^-1^. During a one-hour period of ‘dawn’ and ‘dusk’, light and temperature changes were ramped on a gradient. Humidity was ambient. After ten days of growth, seedlings were transplanted to one-litre pots in standard peat-based compost.

Postharvest (post sample **H**), leaves were stored for two days in a cold store (4 °C) ^15^. Samples for **D0** and **D7** were washed individually in mildly chlorinated water (sodium hypochlorite; 30 ppm ^41^) for two minutes, then rinsed for one minute with distilled water (all at >14 °C to avoid cold-shock). Leaves were dried of excess moisture for one minute using a kitchen salad spinner, then placed in fresh bags, sealed, and stored overnight at 4 °C. Shelf life leaves were stored in the cold and dark (4 °C) for seven days (**D7**) – typical of the use-by date of commercially bagged leaves.

All samples were taken between the hours of 1 – 3 pm to mitigate diurnal fluctuations in phytochemical content and gene expression ^42^. Immediately after each of the aforementioned samples was taken, leaves were frozen using liquid nitrogen and ground into a fine powder using a pestle and mortar. Samples were stored at −80 °C in tubes and lyophilized prior to chemical analysis. A subset of non-lyophilized sample was kept aside for RNA extractions.

### RNA extraction and quality control

RNA for RNAseq and qRT-PCR analyses was extracted using RNeasy Plant Mini Kit (Qiagen, Manchester, UK) according to the manufacturer ‘Plants and Fungi’ procedure. As part of the protocol, an on-column DNase digestion was incorporated according to the RNase-Free DNase Set (Qiagen) protocol. Samples were checked for degradation and contamination prior to sequencing using agarose gel electrophoresis (1%, TAE buffer), Qubit, and NanoPhotometer (Implen, CA, USA) methods. Briefly, ≥2 µg of total RNA was obtained for each sample at a minimum concentration of ≥50 ng.µL^-1^. RNA integrity was determined and evaluated using an Agilent 2100 Bioanalyzer ^43^.

### RNAseq library preparation and sequencing

After QC procedures, sequencing libraries were prepared using NEBNext Ultra RNA Library Prep Kit for Illumina (NEB, MA, USA) following the manufacturer’s instructions, and index codes were added to attribute sequences to each sample. mRNA was purified from total RNA by using poly-T oligo-attached magnetic beads. Fragmentation was carried out using divalent cations under elevated temperature in NEBNext First Strand Synthesis Reaction Buffer (5x). First strand cDNA was synthesized using random hexamer primer and M-MuLV Reverse Transcriptase (RNase H-). Second strand cDNA synthesis was subsequently performed using DNA Polymerase I and RNase H. Remaining overhangs were converted into blunt ends via exonuclease/polymerase activities. After adenylation of 3’ ends of DNA fragments, NEBNext Adaptor with hairpin loop structure were ligated to prepare for hybridization. In order to select cDNA fragments of 150 – 200 bp in length preferentially, the library fragments were purified with an AMPure XP system (Beckman Coulter, MA, USA). 3 µl USER Enzyme (NEB) was used with size-selected, adaptor-ligated cDNA at 37 °C for 15 min, followed by 5 min at 95 °C before PCR. PCR was performed with Phusion High-Fidelity DNA polymerase, Universal PCR primers and Index (X) Primer. Finally, PCR products were purified (AMPure XP system) and library quality was assessed on using an Agilent 2100 Bioanalyzer system.

The clustering of the index-coded samples was performed on a cBot Cluster Generation System using HiSeq PE Cluster Kit cBot-HS (Illumina) according to the manufacturer’s instructions. After cluster generation, the library preparations were sequenced on an Illumina Hiseq platform and 125 bp/150 bp paired-end reads were generated.

### RNAseq bioinformatics

Raw data (raw reads) of FASTQ format were firstly processed through Novogene Co. Ltd. perl scripts. Clean reads were obtained by removing reads containing adapter, reads containing ploy-N, and low quality reads from the raw data. At the same time, Q20, Q30 and GC content of the clean data were calculated.

An index of the reference genome was built using Bowtie (v2.2.3), and PE clean reads were aligned to the reference genome using TopHat (v2.0.12) ^44–46^. TopHat was selected as the mapping tool as it can generate a database of splice junctions based on the gene model annotation file and thus a better mapping result is achieved than other non-splice mapping tools.

HTSeq (v0.6.1) was used to count the read numbers mapped to each gene ^47^. FPKM (Fragments Per Kilobase of transcript sequence per Millions base pairs sequenced) of each gene was calculated based on the length of the gene and reads count mapped to each gene. Differential expression analysis of each sample point/inbred line (three biological replicates) was performed using the DESeq R package (1.18.0) ^46, 48^. The resulting *P*-values were adjusted using Benjamini and Hochberg’s approach for controlling the false discovery rate. Genes with an adjusted *P*-value (<0.05) were assigned as being significantly differentially expressed.

### RNAseq validation by qRT-PCR

Independent RNA extractions were conducted for qRT-PCR validation, and quality checked according to the same protocols and instrumentation as for RNAseq. cDNA synthesis was conducted using qPCRBIO cDNA Synthesis Kit (PCR Biosystems Ltd., London, UK) according to the manufacturer instructions. cDNA was then diluted 10x prior to analysis. All 54 biological samples were tested in triplicate.

PCR primers were designed using PRIMER3 (http://bioinfo.ut.ee/primer3/) using default settings. Ten genes related to GSL biosynthesis and transcription were selected at random for the validation analysis (*BCAT4, CYP83B1, MYB122-1a, MYB51a, SOT16, SUR1, TGG1b, TGG1d, TGG1j,* and *UGT74B1*), with *ACT11* used as a reference gene ^49^. Gene sequences of *E. sativa* were obtained using NovoFinder (Novogene Co. Ltd.), and primer annealing sites were designed to span intron-intron boundaries where possible (see Supplementary Table S5).

Analysis was performed using the 2^-ΔΔCt^ method ^50^ on a Roche LightCycler 480 Instrument and the Advanced Relative Quantification protocol (v1.5.1). Primer efficiencies were determined by analyzing each primer set with log-fold dilutions of cDNA (Supplementary Table S5). 2x qPCRBIO SyGreen Blue Mix Lo-ROX (PCR Biosystems Ltd.) was used to prepare a master mix for all reactions. Reaction volumes totaled 10 µL, and the PCR method used was as per the manufacturer recommendations.

Data were normalized and expressed as the log2-fold change relative to *ACT11*. RNAseq data for each of the tested genes were similarly converted for direct comparison of the two methodologies (Supplementary Figure S3). An ANOVA test found no significant differences between the two data sets for each of the respective genes.

### Intact glucosinolate extraction and analysis by LC-MS

Intact GSLs were extracted according to the protocol used by Bell et al. ^8^. Immediately before LC-MS analysis, samples were diluted with 4 mL of HPLC-grade water. Samples were analyzed in a random sequence with standards and QC samples. External standards of sinigrin (SIN; >99% TLC), GRA (99.86%, HPLC), glucoalyssin (GAL) (98.8%, HPLC), 4OHB (96.19%, HPLC) and GER (99.68%, HPLC) were prepared for quantification of GSL compounds. SIN was used to quantify DGTB, GSV, and DMB, as no standards are available for these compounds. 4OHB was used to quantify the indole GSLs 4MOB and neoglucobrassicin (NGB). All standards with the exception of SIN (Sigma Merck, Gillingham, UK) were purchased from PhytoPlan (Heidelberg, Germany). Recovery of extracted GSLs was calculated by spiking six random samples (in duplicate) with SIN (0.06 µM). The average recovery was 104.8%, indicating excellent preservation of GSLs throughout the extraction process. Limits of detection (LOD) and quantification (LOQ) were established for the method by running serial dilutions of SIN (LOD = 2.14 mg.kg^-1^; LOQ = 6.48 mg.kg^-1^).

LC-MS analysis was performed in the negative ion mode on an Agilent 1260 Infinity Series LC system (Agilent, Stockport, UK) equipped with a binary pump, degasser, auto-sampler, column heater and diode array detector, coupled to an Agilent 6120 Series single quadrupole mass spectrometer. Separation of samples was achieved on a Gemini 3 µm C18 110Å (150 x 4.6 mm) column (with Security Guard column, C18; 4mm x 3mm; Phenomenex, Macclesfield, UK). GSLs were separated during a 40 min chromatographic run, with a 5 min post-run sequence. Mobile phases consisted of ammonium formate (0.1%; A) and acetonitrile (B) with the following gradient timetable: (i) 0 min (A-B, 95:5, v/v); (ii) 0-13 mins (A-B, 95:5, v/v); (iii) 13-22 mins (A-B, 40:60, v/v); (iv) 22-30 mins (A-B, 40:60, v/v); 30-35 mins (A-B, 95:5, v/v); (v) 35-40 mins (A-B, 95:5, v/v). The flow rate was optimized for the system at 0.4 mL min^-1^, with a column temperature of 30 °C, and 20 µl of sample injected into the system. Quantification was conducted using a diode array detector at a wavelength of 229 nm.

MS analysis settings were as follows: Atmospheric pressure electrospray ionization was carried out in negative ion mode (scan range m/z 100–1500 Da). Nebulizer pressure was set at 50 psi, gas-drying temperature at 350 °C, and capillary voltage at 2,000 V. Compounds were identified using their primary ion mass [M-H]-, and comparison to authentic standards ^51, 52^. Data were analyzed using Agilent OpenLAB CDS ChemStation Edition for LC-MS (vA.02.10). GSL concentrations from each time point were averaged over three biological replicates with two technical replicates of each (*n* = 6). This approach was also conducted for GHP and monosaccharide content.

### Glucosinolate hydrolysis product extraction and analysis by GC-MS

GHPs were extracted according to the protocol presented by Ku et al. ^53^ with the following modification: samples were hydrolysed in d.H2O for three hours at 30 °C before extraction with dichloromethane (DCM) for 21 hours. This duration was optimized for maximum yields of GHPs by comparison of extractions for three hours incubation at 30 °C with immediate DCM extraction, and three, nine, and 21 hours post incubation with DCM. GC-MS analysis and GHP identification was conducted according to the method presented by Bell et al. ^15^. Concentrations of all GHPs were calculated as equivalents of SF standard (Sigma).

### Monosaccharide extraction and analysis by HPLC

Free monosaccharides were extracted according to the method presented by Bell et al. ^5^, with the exception that 0.2 g of lyophilized leaf powder was extracted. Extracts were analyzed on an Agilent 1100 series HPLC system equipped with a binary pump, degasser, and auto-sampler, with an external column heater (50 °C). A Bio-Rad Aminex HPX-87H (300 x 7.8 mm, 9 μm particle size) column with a Micro-Guard Cation H guard column (Bio-Rad, Watford, UK) was used to achieve separation with an isocratic gradient of 5 mM sulfuric acid, and a flow rate of 0.6 mL per min. A Polymer Laboratories ERC-7515 refractive index detector (Church Stretton, UK) was used to detect monosaccharides. Compounds were quantified using authentic standards and analyzed with Agilent ChemStation software (Santa Clara, CA, USA).

### Sulfur content analysis by ICP-OES

Lyophilized samples were weighed into acid washed glass boiling tubes, pre-digested in 70% nitric acid for 24 hours, before being heated to 90 °C for two hours using a heat block. Once cooled, these were filtered through a 0.45 µM syringe filter, and diluted give an acid concentration of 3%. These samples were analysed using inductively coupled plasma-optical emission spectroscopy (ICP-OES) (Perkin Elmer Optima 7300 DV). Sulphur content was determined using the radial signal at 181.975 nm.

### Statistical analyses

All statistical analyses (not included in bioinformatics sections) were performed using XL Stat (Addinsoft, Paris, France). Shapiro-Wilk normality tests were conducted for all variables and fit a normal distribution. ANOVA with post-hoc Tukey’s Honest Significant Difference (HSD) tests were performed to generate multiple pairwise comparisons between sampling points for each cultivar (i.e. **H** vs. **D7** for cultivar **B**) and between cultivars at each respective time point (i.e. **A** vs. **B** for time point **H**) for phytochemical data (Supplementary Data File S1). All PCAs were performed using Pearson correlation coefficient analysis, *n*-1 standardization, Varimax rotation, and Kaiser Normalization. Phytochemical data were regressed onto the gene expression data as supplementary variables for the targeted analysis.

## Supplementary Materials

Fig. S1. RNAseq experimental design and sampling diagram.

Fig. S2. RNAseq sample replicate Pearson correlation analysis.

Fig. S3. RNAseq gene expression validation by qRT-PCR.

Table S1. Summary of genome assembly and annotation statistics.

Table S2. Reference genome sequence composition.

Table S3. Predicted protein-coding genes within the *E. sativa* reference genome.

Table S4. Numbers of genes with homology or functional assignment within the *E. sativa* genome annotation.

Table S5. qRT-PCR primer sequences and amplification efficiencies.

Data file S1. Phytochemical Analyses of Variance (ANOVA) pairwise comparisons and RNAseq gene expression correlation analysis.

Data file S2. Sulfur metabolism and glucosinolate biosynthesis gene expression values and annotation descriptions.

## Acknowledgments

### General

The authors would like to thank: the Vegetable Plant Breeding Team at Elsoms Seeds Ltd.; members of the Genomics Pipelines Group in the BBSRC National Capability in Genomics and Single Cell (BB/CCG1720/1) at Earlham Institute; Dr. Yunan Lin and Irene Wei for project and technical support for genome annotation, RNAseq, and bioinformatics at Novogene Co. Ltd.; Matthew Richardson for maintenance of controlled environment facilities at the University of Reading Controlled Environment Laboratory; and Dr. Marcia Boura for advice on qRT-PCR.

### Funding

Dr. Luke Bell was supported by a BBSRC Case Award (BB/J012629/1) in partnership with Elsoms Seeds Ltd. (Spalding, UK) and Bakkavor Group Ltd. (Spalding, UK) for *de novo* genome sequencing and assembly. Dr. Luke Bell, Dr. Martin Chadwick, and Manik Puranik were supported by a BBSRC LINK award (BB/N01894X/1) for all other work.

### Author contributions

LB and CW conceived and designed the experiment and analyses. RT and SK produced the breeding line seed material for genome sequencing and the RNAseq experiment. LB grew plants in controlled environment, performed RNA extractions and quality control, qRT-PCR validation, glucosinolate analysis by LC-MS, and hydrolysis product analysis by GC-MS. MP performed sugar analysis by HPLC. MC performed sulfur content analysis by ICP-OES. LB performed ANOVAs, Pearson’s correlation analyses, and Principal Component Analyses of all phytochemical and gene expression data. LB wrote the paper, with contributions from MC, RT, SK, and CW. Funding was obtained by LB, LM, and CW.

### Competing interests

The authors declare no conflicts of interest.

### Data and materials availability

Full reference genome sequence and annotation can be found in the European Nucleotide Archive. Assembly accession no: GCA_902460325; Study ID: PRJEB34051; Sample ID: ERS3673677; Annotation accession number ERZ1066251.

## Supplementary Materials

**Table S1.**
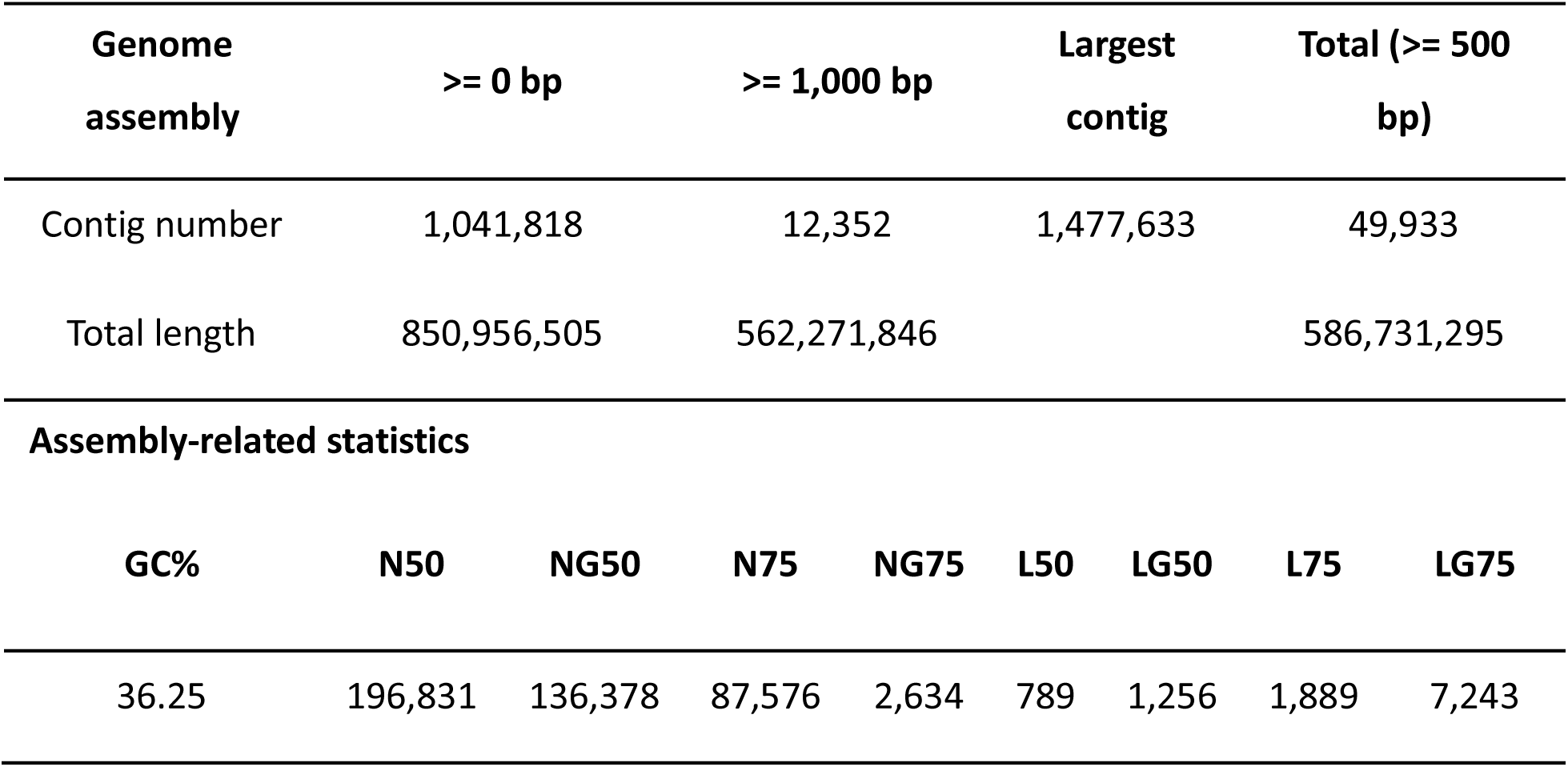
Summary of genome assembly and annotation statistics.

**Table S2.** Transposable elements content in the reference genome.

| Type | Denovo + Repbase* | | TE proteins<sup>\$</sup> | | Combined TEs<sup>£</sup> | |
| --- | --- | --- | --- | --- | --- | --- |
|  | Length<br>(bp) | % in<br>genome | Length<br>(bp) | % in<br>genome | Length<br>(bp) | % in<br>genome |
| DNA | 69,251,054 | 8.14 | 21,607,510 | 2.54 | 76,517,426 | 8.99 |
| LINE | 20,290,781 | 2.38 | 17,153,783 | 2.02 | 28,200,567 | 3.31 |
| SINE | 2,134,305 | 0.25 | 0 | 0 | 2,134,305 | 0.25 |
| LTR | 311,377,915 | 36.59 | 91,001,347 | 10.69 | 317,124,290 | 37.27 |
| Other<sup>^</sup> | 106,176 | 0.01 | 0 | 0 | 106,176 | 0.01 |
| Unknown<sup>^^</sup> | 155,033,031 | 18.22 | 0 | 0 | 155,033,031 | 18.22 |
| Total | 547,675,259 | 64.36 | 129,571,414 | 15.23 | 563,873,839 | 66.26 |
| * = RepeatMasker based on the uclust algorithm combined with the known Repbase and *de novo* repeat library created by RepeatModeler / Repeat Scout / LTR_finder |  |  |  |  |  |  |
| <sup>\$</sup> = RepeatProteinMask based on Repbase | | | | | | |
| <sup>£</sup> = The non-redundant set of results combining Denovo+Repbase TEs and TE proteins |  |  |  |  |  |  |
| <sup>^</sup> = Repeats that can be classified by RepeatMasker, but not included by classes above |  |  |  |  |  |  |
| <sup>^^</sup> = Repeats that could not be classified by RepeatMasker |  |  |  |  |  |  |

**Table S3.** Predicted protein-coding genes within the *E. sativa* reference genome.

| Gene set |  | Number | Average gene length (bp) | Average CDS length (bp) | Average exons per gene | Average exon length (bp) | Average intron length (bp) |
| --- | --- | --- | --- | --- | --- | --- | --- |
| *De novo* * | Augustus | 50,179 | 1,701.56 | 1,024.89 | 4.48 | 228.88 | 194.57 |
|  | Glimmer HMM | 73,989 | 1,335.36 | 725.53 | 3.03 | 239.16 | 299.86 |
|  | SNAP | 80,264 | 1,231.13 | 728.3 | 4.02 | 180.98 | 166.27 |
|  | Geneid | 100,165 | 2,127.25 | 585.82 | 3.07 | 191.03 | 745.87 |
|  | Genscan | 71,813 | 3,942.04 | 785.32 | 3.92 | 200.11 | 1,079.4 |
| Homolog^ | *Arabidopsis lyrata* | 32,667 | 1,867.18 | 1,084.09 | 4.86 | 223.12 | 202.93 |
|  | *Arabidopsis thaliana* | 27,416 | 1,870.34 | 1,218.4 | 5.13 | 237.58 | 157.91 |
|  | *Brassica napus* | 101,040 | 1,764.75 | 1,001.16 | 4.91 | 204.06 | 195.48 |
|  | *Boechera stricta* | 27,416 | 2,006.68 | 1,181.2 | 5.09 | 231.86 | 201.61 |
|  | *Capsella rubella* | 26,521 | 1,958.82 | 1,248.6 | 5.19 | 240.53 | 169.46 |
|  | *Raphanus sativus* | 49,733 | 2,064.75 | 1,194.41 | 4.94 | 241.57 | 220.66 |
| RNAseq^^ | Cufflinks | 43,200 | 2,848.16 | 1,723.58 | 6.03 | 285.65 | 223.41 |
|  | PASA | 37,870 | 1,744.08 | 1,034.72 | 4.77 | 216.9 | 188.13 |
| EVM |  | 59,643 | 1,665.12 | 926.2 | 4.17 | 222.2 | 233.23 |
| PASA-update |  | 59,491 | 1,656.25 | 929.33 | 4.17 | 222.9 | 229.36 |
| **Final set** |  | **45,438** | **1,889.6** | **1,069.44** | **4.76** | **224.81** | **218.3** |

\* = The combined results by EVM of *5-ab initio* gene predictions
^ = The combined results by EVM of homology-based gene prediction
^^ = The combined results by EVM of transcriptome data sets

**Table S4.** Numbers of genes with homology or functional assignment.

| Database |  | Number of annotated genes | Gene annotation % |
| --- | --- | --- | --- |
| NR |  | 44,630 | 98.2 |
| Swiss-Prot |  | 35,309 | 77.7 |
| KEGG |  | 32,589 | 71.7 |
| InterPro | All | 37,067 | 81.6 |
|  | Pfam | 34,541 | 76 |
|  | GO | 25,820 | 56.8 |
| Annotated |  | 44,655 | 98.3 |
| Total |  | 45,438 | - |

**Table S5.**
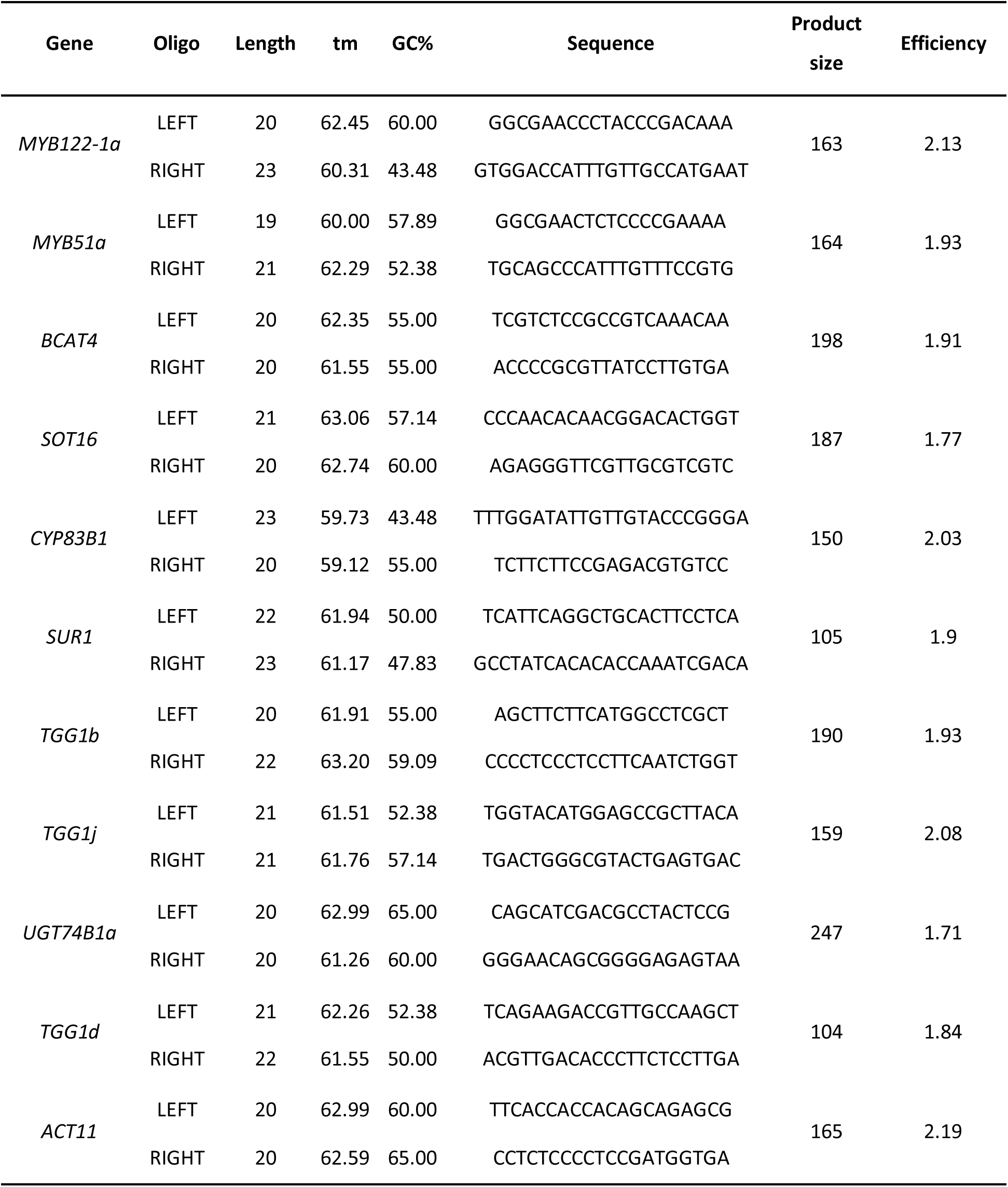
qRT-PCR primers and efficiencies.

**Supplementary Figure S1.**
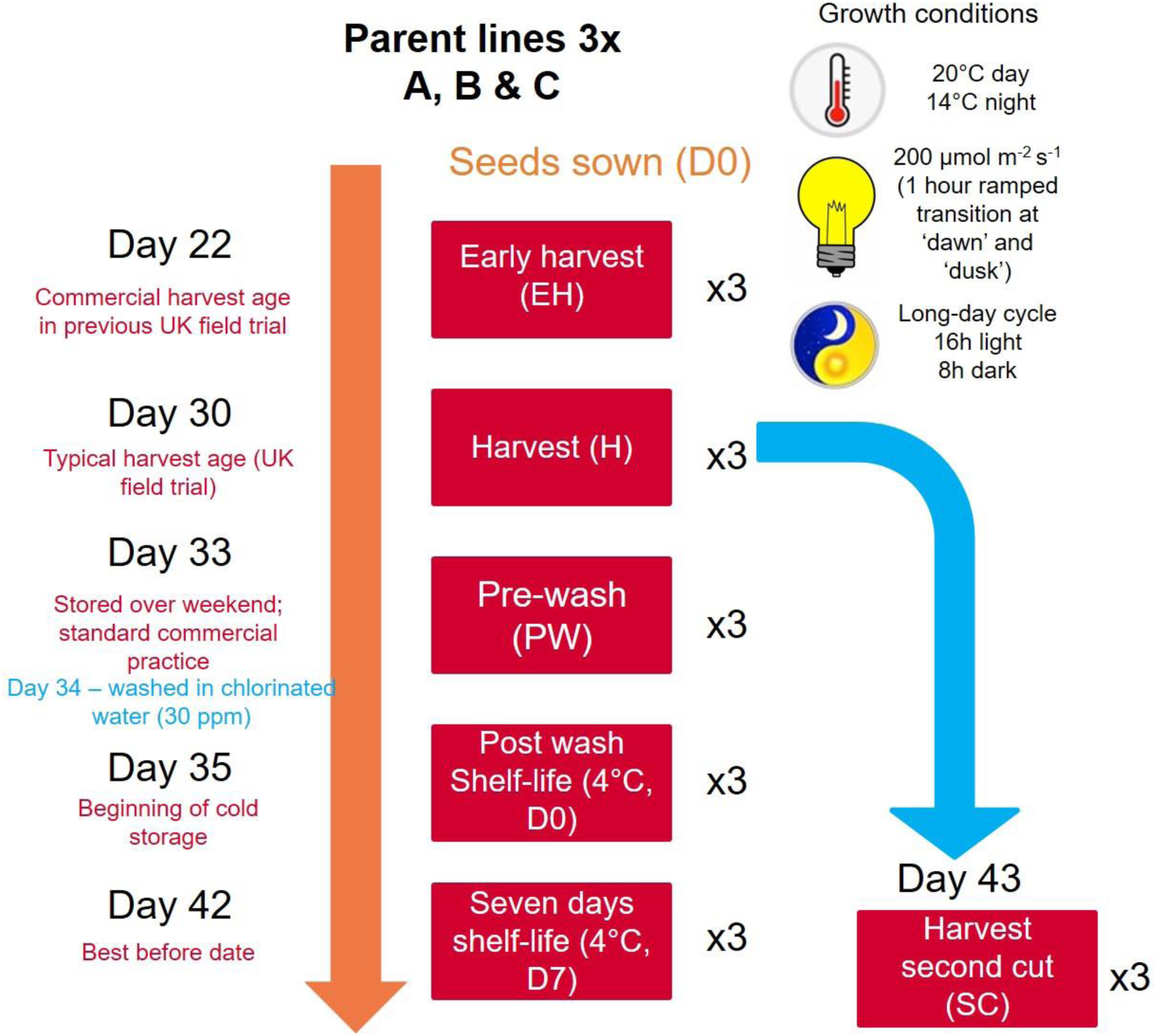
RNAseq experimental design and sampling diagram. Three elite inbred lines of *Eruca sativa* were grown under controlled environment conditions and sampled at each of the six time points indicated (in triplicate). Abbreviations: early harvest (**EH**), harvest (**H**), second harvest (**SC**), pre-wash (**PW**), post-wash (**D0**), and seven-day shelf life (**D7**).

**Supplementary Figure S2.**
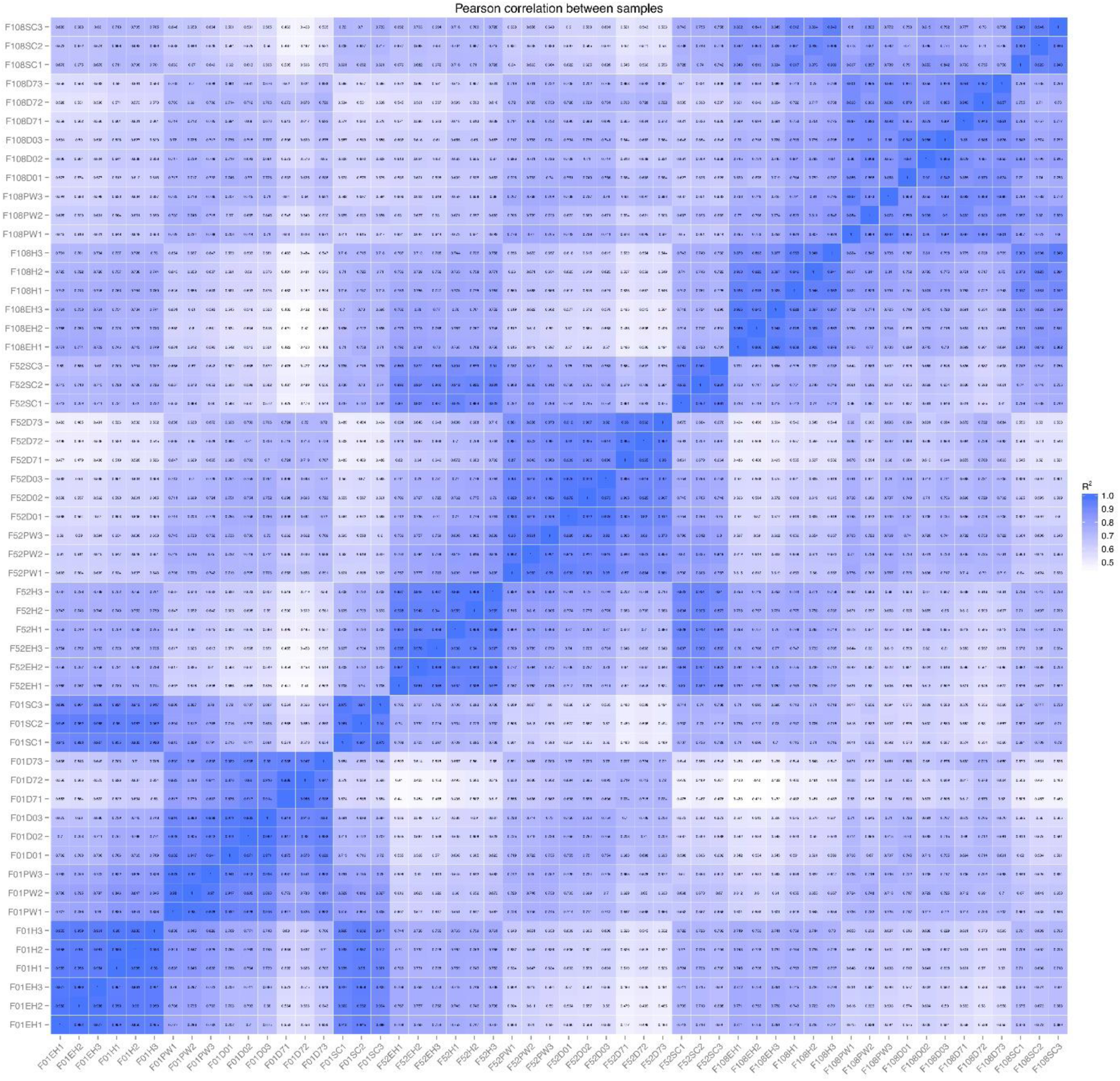
Pearson correlation matrix of RNAseq biological sample replicate gene expression values. Replicates of each sample showed a high degree of correlation (*r*^2^ = >0.884) indicating robust reproducibility of gene expression between the individual plants tested at each respective sample point.

**Supplementary Figure S3.**
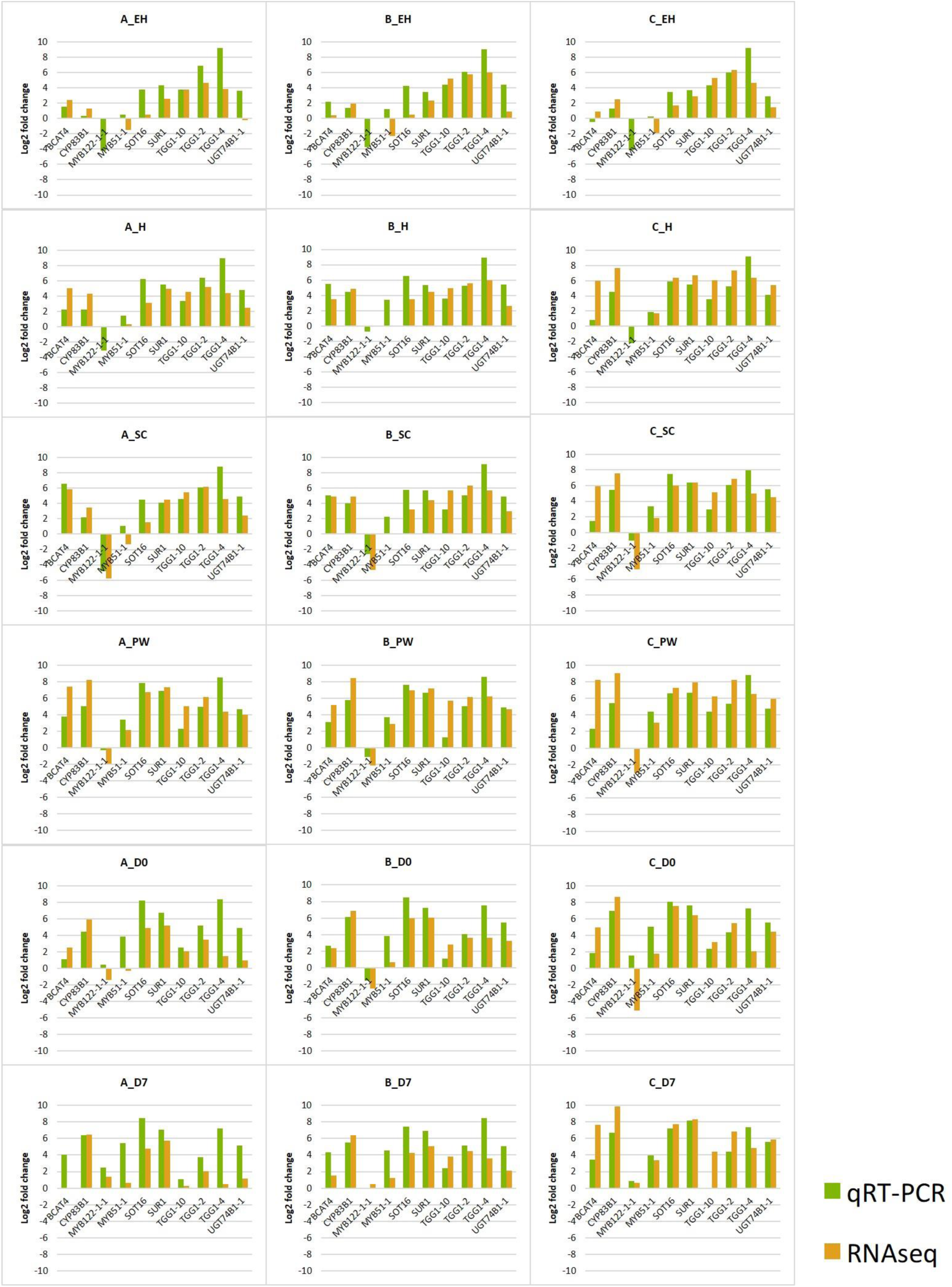
qRT-PCR (green) vs. RNAseq (orange) gene expression of ten randomly selected glucosinolate biosynthesis and hydrolysis-related genes. Data are expressed as the normalized log2 fold-change in expression relative to the reference gene *ACT11*. ANOVA revealed no significant difference between the two data sets. Abbreviations: early harvest (**EH**), harvest (**H**), second harvest (**SC**), pre-wash (**PW**), post-wash (**D0**), and seven-day shelf life (**D7**).

**Supplementary Data File S1**

Analysis of Variance outputs with Tukey’s HSD pairwise comparisons between sample points and each respective rocket breeding line: Tab 1 – glucosinolate analysis; Tab 2 – glucosinolate hydrolysis product analysis; Tab 3 – sugar analysis. Highlighted values are significant at the following levels: *P* = <0.05 (yellow), *P* = 0.01 (orange), and *P* = 0.001 (green). Tab 4 contains a Pearson’s correlation analysis matrix for sulfur and glucosinolate-related gene expression values and phytochemical observations. Values in bold are significant correlations at the *P* = 0.001 threshold.

**Supplementary Data File S2**

RNAseq read counts, log2-fold changes, *P*-values, and adjusted *P*-values (padj) for sulfur metabolism, glucosinolate biosynthesis, hydrolysis, and transport genes for each of the three rocket lines and the respective sample points. Significant up/down regulation is denoted by green/red highlighting, respectively. KEGG annotation numbers and UniProt gene descriptions for orthologous genes in *Arabidopsis thaliana* are provided.

